# Mechanistic and machine learning models design human gut consortia that robustly inhibit *Clostridioides difficile*

**DOI:** 10.64898/2026.08.04.742850

**Authors:** Hanhyeok Im, Yiyi Liu, Jaron Thompson, Gabriel I. Miranda-Contreras, Kaixiwang Loke, Rebecca Bacon, Lauren Lucas, Daniel Amador-Noguez, Simon M. Gray, Ophelia S. Venturelli

## Abstract

Identifying design principles for robust inhibition of human pathogens is a major goal of microbiome engineering. By building synthetic microbial communities from the bottom-up guided by Bayesian active learning, we investigate the *Clostridioides difficile* growth landscape across thousands of species-metabolite conditions. Mechanistic consumer resource and machine learning models uncover significant interactions linking metabolites and species, and exhibit concordance with microbial interactions identified in human microbiome datasets. Guided by machine learning and mechanistic models, we elucidate microbial communities capable of robustly inhibiting *C. difficile* across diverse nutrient environments *in vitro*. Metabolomic profiling identified sorbitol, mannitol, and proline as key metabolites mediating community-driven *C. difficile* inhibition. Model-optimized communities significantly reduced *C. difficile* colonization in the murine gut whereas a model-designed non-inhibitory community did not exhibit this property. Communities display consistent metabolomic patterns *in vitro* and *in vivo*, suggesting that resource competition is a design principle of robust inhibition. Together, these findings establish an integrated experimental and computational framework for the rational design of microbial communities that robustly suppress pathogens across diverse environmental contexts.

## INTRODUCTION

The human gut microbiome shifts dynamically between a healthy and a dysbiotic state due to the tug-of-war between commensals and pathogens^1,2^. *Clostridioides difficile (C. difficile* or CD) is an opportunistic bacterial pathogen that can infect the human gastrointestinal tract, leading to *C. difficile* infection (CDI)^3,4^. Antibiotics, the standard first-line treatment for CDI, are often associated with recurrent infections (rCDI) due to their disruption of the gut microbiome^5,6^. While fecal microbiota transplantation (FMT) has shown efficacy in treating rCDI, its reliance on donor microbiota introduces challenges such as inconsistent patient outcomes and potential transfer of antibiotic-resistant organisms^7–9^. Overcoming these challenges, certain defined microbial consortia, such as VE303, have been evaluated in clinical trials for CDI treatment over the past decade, while donor-derived microbiota-based therapeutics have gained FDA approval^10–13^. These consortia are often identified via screening of reduced complexity communities with the desired functions from healthy donors as opposed to being designed based on mechanistic insights^14^. Our lack of understanding of the mechanisms of action or failure modes of bacterial therapeutics limits our ability to apply these insights more broadly to other pathogens, microbiomes or disease contexts.

Resource competition is a key mechanism that shapes ecological dynamics and microbial colonization of the human gut^15,16^. Therefore, this mechanism modality has potential for impacting pathogens such as *C. difficile*^17,18^. Several metabolites enhance *C. difficile* colonization and toxicity^19^, including amino acids^18,20^, mucus-derived sugars^21^, sorbitol^22^, succinate^23^, trehalose^24^, mannitol^25,26^, and cellobiose^27^. As a proof of concept, a five-member consortium of sialic acid and *N*-acetylglucosamine utilizers reduced *C. difficile* colonization in antibiotic-treated mice via competition for mucosal sugars^28^. Corroborating the role of resource competition influencing *C. difficile* disease severity, an avirulent strain of *C. difficile* protected mice from virulent *C. difficile*-induced colitis via amino acid depletion^29^. Consistent with these results, non-toxigenic *C. difficile* strains protected hamsters from infection^30^. This implies that resource competition has potential as a design rule for bacterial therapeutics that inhibit *C. difficile*.

Top-down community assembly approaches that reduce the complexity of a microbiome have been applied to pathogen suppression with variable success^17,31^. In contrast, bottom-up synthetic community assembly coupled with computational modeling can be leveraged to explore combinatorial landscapes of species and environmental factors, generating information-rich datasets that elucidate mechanistic insights and map initial species composition to pathogen abundance^32,33^. Considering variability in species composition and nutrient environments is critical to understanding *C. difficile* colonization due to substantial inter-individual variability and intra-individual dietary shifts and other environmental factors^34,35^. Further, *C. difficile* may exploit different metabolic niches over time due to its metabolic plasticity^36,37^. For example, the consumer resource model (CR), a dynamic mechanistic model that captures metabolite-mediated interactions^38^, has been used to elucidate the role of resource competition in driving microbial community assembly^39,40^. Since the design spaces grow exponentially with the number of input variables, Bayesian experimental design that seeks to maximize information gain from limited experiments^41,42^ can enable efficient exploration of the large species and resource design space.

Mechanistic models are based on known interaction modes shaping community dynamics. By contrast, machine learning methods can learn to capture complex relationships between species and metabolites from data^43^. Among machine learning methods, models based on the construction of an ensemble of regression trees have been shown to excel at the prediction of microbial community data^44–47^. For example, Gradient Boosting (GB) is a tree-based model that displays improved prediction performance with skewed or heavily tailed feature distributions, a common feature of microbiome datasets, compared to other machine learning models such as neural networks^47–49^. To uncover interactions shaping microbial communities, interpretable artificial intelligence techniques such as SHapley Additive exPlanation (SHAP), applied to the trained models, can elucidate the contributions of input variables on the model’s output prediction^50–52^.

Leveraging high-throughput bottom-up synthetic microbial community assembly and model-guided experimental design, we investigated *C. difficile* growth in human gut communities across different nutrient environments. We constructed a species design space of 29 human gut commensals and a resource design space of 11 metabolites critical for *C. difficile*’s metabolic niche in the mammalian gut^20–27^. To efficiently explore the large design space (2^40^ considering presence/absence of variables, ∼550 billion), we used design-test-learn (DTL) cycles coupled with a CR model and Bayesian experimental design to generate an information-rich dataset of 2,743 unique monoculture and community conditions. To evaluate resource competition as a design rule for *C. difficile* inhibition, we analyzed differences in inferred interactions and model prediction performance using a flexible GB model and a mechanistic CR model. Communities optimized for *C. difficile* inhibition or non-inhibition across a wide range of nutrient environments were selected for validation *in vitro* and in a murine model. Our results demonstrate that both models identified communities that robustly inhibited *C. difficile* growth across nutrient environments *in vitro*. In a murine model, model-designed communities for *C. difficile* inhibition significantly reduced *C. difficile* abundance in the mammalian gut, whereas a model-designed non-inhibitory community displayed no-impact on *C. difficile*’s colonization *in vivo*. In sum, our framework integrates high-throughput bottom-up community assembly, mechanistic modeling and ML to establish foundation for the discovery of design principles of community-level interactions impacting human pathogens.

## RESULTS

### Investigating the C. difficile growth landscape using Bayesian experimental design

To investigate *C. difficile* inhibition by human gut commensals across diverse nutrient environments, we selected a set of diverse human gut species and metabolites that have been previously shown to be used by *C. difficile.* The species design space consisted of 28 commensal species and *C. difficile* **(Figure 1A)** and included representative members of major phyla in the human gut microbiome (Proteobacteria, Bacteroidetes, Firmicutes, and Actinobacteriota). The species in this collection are metabolically diverse^53^ and most are highly prevalent across individuals^54,55^, thus representing a healthy human gut microbiome. Although the 29-species are a sub-system of the human gut microbiome, the community recapitulates much of the metabolic capacity of large synthetic communities designed to represent the full complex microbiome (hCom1 and hCom2)^56^. Its distribution of pairwise functional distances between member species is comparable to that of hCom1 **(Figure S1B)**, indicating that members are not functionally clustered, and it captures ∼77% of the KEGG Ortholog (KO) metabolic pathways of the 119-species hCom2 **(Figure S1C)**, together supporting its suitability as a representative model system. Many species in the commensal design space have previously established links to *C. difficile* inhibition. For example, *Peptacetobacter hiranonis* (formerly *Clostridium hiranonis*, PH) and *Lachnoclostridium scindens* (formerly *Clostridium scindens*, LS) were shown to inhibit *C. difficile in vitro* and ameliorate disease severity in a murine model^31,32,42^. The resource design space consists of 11 resources previously shown to mediate *C. difficile* colonization of the murine gut or virulence activities. These resources include amino acids^20^, mucus-derived sugars (acetyl-neuraminic acid or sialic acid, acetyl-glucosamine or GlcNAc, acetyl-galactosamine or GalNAc, galactose, and mannose)^21^, sorbitol^22^, succinate^23^, trehalose^24^, mannitol^25,26^, and cellobiose^27^.

**Figure 1.**
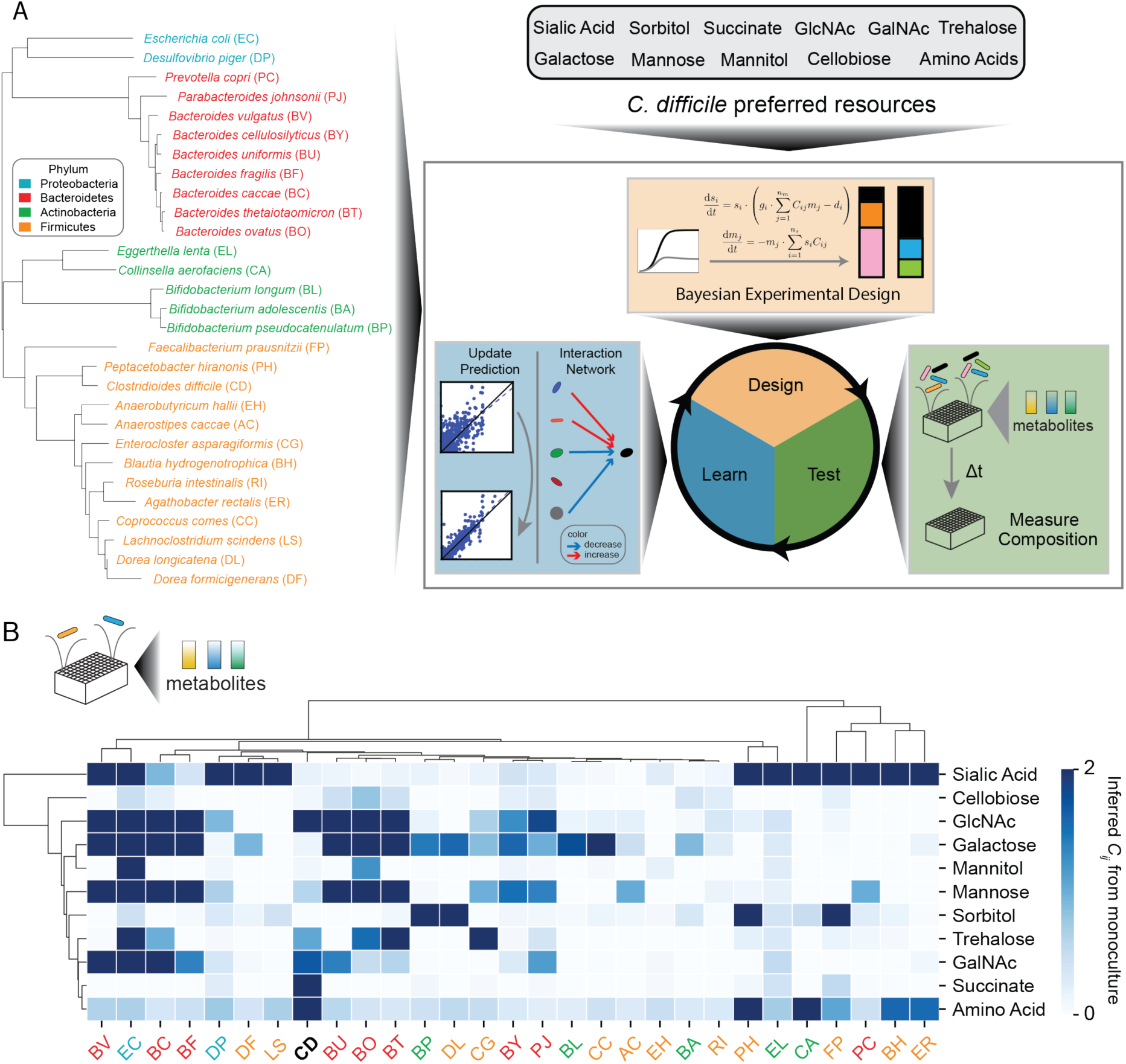
Exploration of a vast species-metabolite design space to identify communities that robustly inhibit *C. difficile* in response to nutrient variations. (A) Overall project schematic of model-guided community design. The design space is comprised of commensal species and *C. difficile* preferred resources. The phylogenetic tree of the species design space is composed of 29 highly prevalent and diverse species, generated by multiple sequence alignment of 37 housekeeping genes using PhyloSift^61^. Species color indicates phylum. Species two-letter abbreviations are noted in brackets next to species names. The resource design space consists of 11 resources, all major substrates previously shown to be used by *C. difficile* during *in vivo* colonization. The design-test-learn (DTL) cycles were applied for sufficient exploration of the species and metabolite design space. Bayesian experimental design is applied on the CR model to identify experiments to constrain model parameter uncertainty. Given a set of n_s_ species and n_m_ resources, the proposed CR model is a set of ordinary differential equations. The DTL cycles terminate when the predictive performance of the CR model saturates. The entire dataset from all DTL cycles were used for model-guided community designs. (B) Clustered heatmap of inferred *C_ij_* parameters in the CR model of monoculture growth dynamics. Species colors represent phylum. Target species *C. difficile* (CD) is bolded and highlighted in black.

Due to the immense size of the ∼550 billion combinatorial design space, we aimed to efficiently explore the species-resource design space by model-guided DTL cycles **(Figure 1A)**. To explore the role of resource competition in shaping *C. difficile* growth, we used a modified CR model capturing resource competition. The rate of change of species abundance is coupled to resource consumption through the coefficient *C_ij_*, the rate at which species *i* consumes resource *j*, together with a species-specific death rate, *d_i_*. We additionally introduce a species-specific growth efficiency parameter *g_i_* that scales how efficiently consumed resources are converted into biomass. In this way, *C_ij_* reflects resource consumption and thus competition among species, while *g_i_* separately accounts for each species’ growth yield. Our framework involves construction of an initial model informed by monoculture and community dynamics in various media conditions. In the second stage, we apply Bayesian experimental design^41^, a technique used to identify informative conditions for experimental characterization that seek to minimize model parameter uncertainty and thus maximize information from limited experiments (**Methods**). To inform the model, we studied monocultures and 2–5 member communities to evaluate low-richness inhibition of *C. difficile*, while keeping the number of species and microbial genes similar across conditions.

In our initial dataset, we characterized the resource utilization capability of each species in our design space in different nutrient environments to provide deeper insights into metabolic niche overlap with *C. difficile* and to inform our mechanistic computational model. Our media conditions contained individual metabolites at different concentrations (1 g/L in **Figure S2A**, 5 g/L in **Figure S2B**, or 10 g/L in **Figure S2C**) to uncover the relationship between species growth and metabolite concentrations. Since amino acids can be used as a major energy source for *C. difficile*^20^, we considered conditions with amino acids in the absence of preferred carbon sources (No Carb condition). Species growth was quantified by computing the area-under-the-curve (AUC) over 24 h based on absorbance at 600 nm (OD_600_) measurements (899 unique monoculture growth conditions) **(Table S5)**. While *C. difficile* monoculture displayed growth in the No Carb condition likely due to Stickland amino acid metabolism, its growth was significantly enhanced in the presence of all single carbon sources **(Figure S2 A-C)**. These data demonstrate that all selected resources enhanced *C. difficile* monoculture growth, consistent with the previous studies^20–27^. Growth of the other species varied with nutrient environment. At 1 g/L, only a subset of species, including *L. scindens* (LS), *P. hiranonis* (PH), *Dorea longicatena* (DL), *Escherichia coli* (EC), and several Bacteroides spp., grew on multiple carbohydrates relative to the No Carb condition **(Figure S2A)**, whereas at higher concentrations most species did so (28 of 29 at 5 g/L, **Figure S2B**; 25 of 29 at 10 g/L, **Figure S2C**). At the highest concentration, growth profiles clustered by phylogeny, indicating that phylogenetic relatedness is informative of functional similarity, with Bacteroides spp. grouping together by their utilization of mucin-derived sugars^57–59^ **(Figure S2C)**.

We fit a CR model to monoculture data (CR-mono) (377 total parameters) to provide insight into the resource utilization capabilities of each species (No Carb and each carbohydrate at three concentration levels). The regularization parameters for minimizing model overfitting (penalizing non-zero parameter values) are estimated using the expectation-maximization algorithm (**Methods**). Resource utilization of resource *j* by species *i* occurs when the mean of *C_ij_* is at least two standard deviations from zero, which for a normal distribution indicates 95% of true *C_ij_* values will be nonzero^60^. Based on this criterion, *C. difficile* uses all resources except sorbitol in monoculture **(Figure 1B)**. Using Jaccard indices of shared resource utilization determined by the *C_ij_* parameters, we quantified the metabolic niche overlap between *C. difficile* and each species. *Bacteroides ovatus* (BO) and *Bacteroides cellulosilyticus* (BY) displayed high predicted niche overlap with *C. difficile*, whereas *Bifidobacterium adolescentis* (BA), *Collinsella aerofaciens* (CA), and *Eubacterium rectale* (ER) exhibited low overlap based on monoculture data **(Figure S2D)**. In sum, *C. difficile* can utilize all resources in the design space, and human gut species displayed distinct resource utilization profiles with varying degrees of overlap with *C. difficile*.

*Bayesian experimental design explores a broad region of the species-metabolite design space.* To reveal the patterns of *C. difficile* growth across the metabolite-species design space, we characterized communities containing *C. difficile* in a wide range of nutrient environments. Specifically, we tested 22 media conditions, including a mixture of all carbohydrates and all carbohydrates except one (1 g/L for each carbohydrate) in media containing a mixture of amino acids at two distinct concentrations (5 g/L or 10 g/L). We constructed 5-member communities including *C. difficile* in each media condition and measured species abundances at two time points (16 and 24 h) **(Table S5)**. The absolute abundance of each species was estimated by computing the product of OD_600_ and relative abundance based on 16S rRNA gene sequencing^32,33,42,62^ (896 unique conditions) **(Figure 2A)**. The abundance of *C. difficile* in 5-member communities was significantly enhanced as the amino acid concentration was increased (5 g/L to 10 g/L) **(Figure S3A),** consistent with the major role of Stickland metabolism in enabling *C. difficile* growth. The removal of single carbohydrates did not impact the *C. difficile* abundance distribution in the context of the mixture **(Figure S3A**). When we sorted *C. difficile* abundance by the constituent species in each community, several species appeared to be associated with lower *C. difficile* levels **(Figure S3B).** Communities containing EC, ER, BC, or RI exhibited median *C. difficile* absolute abundances ranging from 0.09 to 0.13, lower than the highest median value of 0.21 observed for communities containing DL **(Figure S3B)**. However, the broad distribution of *C. difficile* abundance across communities containing these species indicated that outcomes were highly dependent on community composition and nutrient environment. Additionally, species with high monoculture Jaccard index of niche overlap with *C. difficile* (EC, BO, BC, and BT) **(Figure S2D)** displayed highly variable *C. difficile* abundance distributions in the community context **(Figure S3B)**, demonstrating that metabolic niche overlap based on monoculture data may not be sufficient to provide insight into *C. difficile* inhibition in microbial communities.

**Figure 2.**
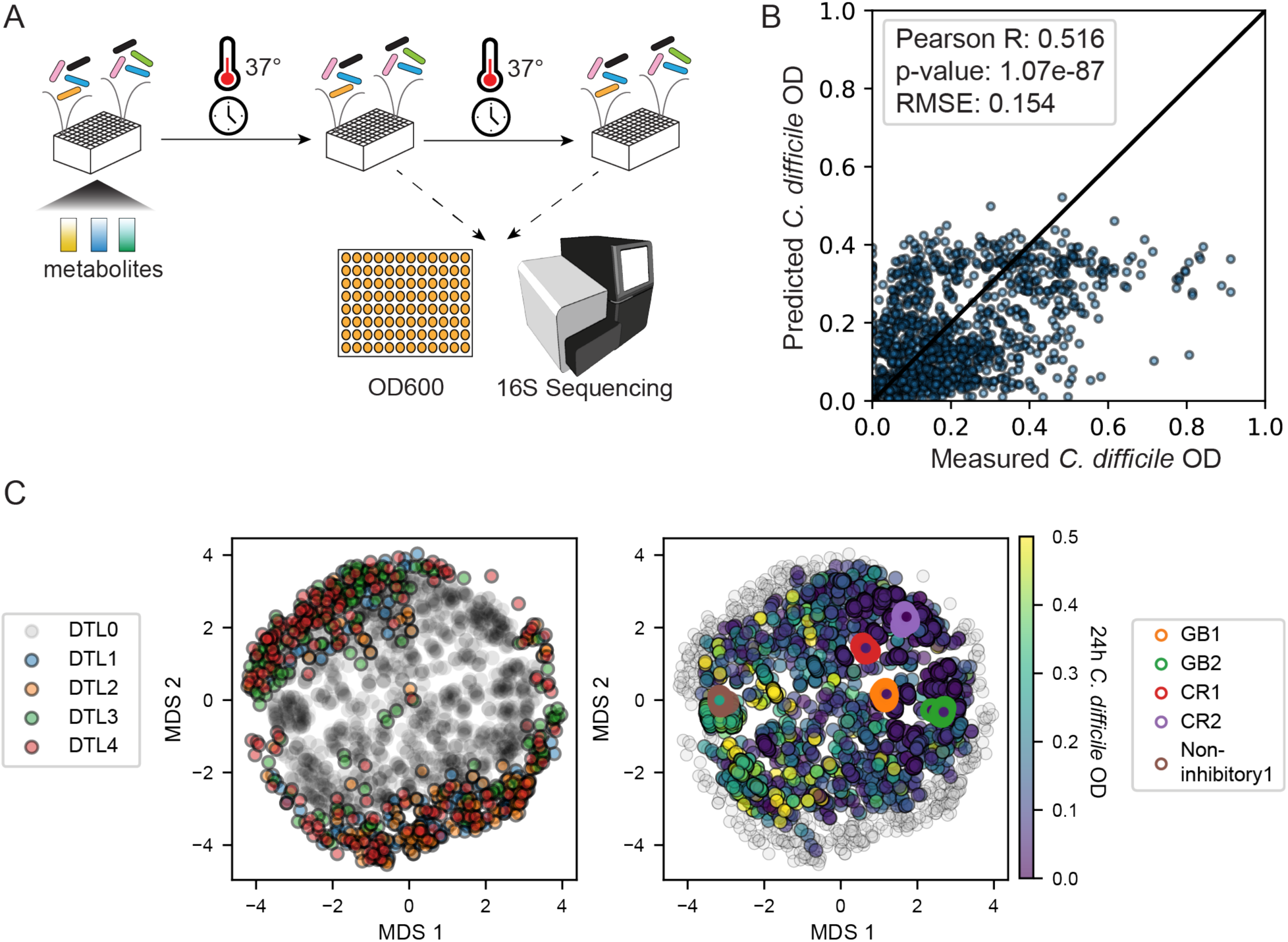
Bayesian model-guided exploration of species-resource landscapes improve model prediction of species abundance and provide biological insights into factors affecting *C. difficile* abundance. (A) The bottom-up community assembly workflow for testing the *C. difficile* inhibition of various species combinations in different metabolite combinations. The species and metabolites tested in each experimental condition are either rationally designed in DTL0 or generated from Bayesian optimization for maximum information gain. Synthetic communities are cultured in 96-well deep well plates in anaerobic conditions and incubated at 37°C. At various timepoints, the communities are sampled for species absolutes abundance, as determined by total cell density (OD_600_) and species relative abundance. Species relative abundances are determined using multiplexed 16S rRNA sequencing. (B) Scatterplot of measured 24 h *C. difficile* OD versus predicted 24 h *C. difficile* OD of CR in 20-fold cross validation. 20-fold cross validation is performed on monoculture and community growth dynamics from all DTL cycles. Prediction performances are evaluated by Pearson correlation and mean squared error between measured and predicted 24 h *C. difficile* OD. Due to the number of unique experimental conditions, individual replicates are omitted from the figure but are available in Table S5. (C) Multidimensional scaling (MDS) projection of the communities with resources design space. Each condition is encoded as a binary vector capturing species phylogeny and metabolite presence/absence, so that proximity in the projection reflects similarity in both phylogenetic composition and nutrient environment. Each point represents a unique experimental condition. Colors represent DTL cycle (left) and 24 h *C. difficile* OD_600_ (right). Marker edge color in the right plot represents communities further validated by *in vitro* exo-metabolomics. GB1 (BF, BU, EC, and RI) and GB2 (BC, CC, EC, and DF), communities optimized by the GB model for low predicted *C. difficile* OD_600_; CR1 (BO, BH, EC, and CA) and CR2 (BO, BT, EC, and FP), community optimized by the CR model for low predicted *C. difficile* OD_600_; Non-inhibitory1 (EH, BH, BL, and EL), non-inhibitory community with high predicted *C. difficile* OD_600_. Only conditions inoculated with *C. difficile* are included in the right plot. Marker with gray edge in the right panel denote conditions not inoculated with *C. difficile*, which therefore lack a *C. difficile* OD600 value.

To construct the initial CR model informed by monoculture and community data, we fit the CR model to DTL0 containing 1,795 unique experimental conditions, including 896 communities and 899 monocultures. For the CR model, parameters that govern the degree of regularization (penalizing non-zero parameter values) are optimized using the expectation-maximization algorithm (**Methods**). This algorithm was chosen to simultaneously estimate species-specific measurement noise variances and perform regularization of hyperparameters via joint inference across monoculture and community data. To further constrain the uncertainty of inferred model parameters, we used Bayesian experimental design to select combinations of species and metabolites that maximize information content as quantified by the expected Kullback-Leibler divergence between the posterior and prior parameter distributions. Briefly, we fit the model using Bayesian inference to approximate the posterior parameter distribution as a Gaussian^41^. The parameter distribution is then used as a prior to guide experimental designs of species and resource combinations **(Figure S4 A and B)**. For computational efficiency, we considered 5-6 carbohydrates out of 10 total for the Bayesian experimental design to reduce the design space. We prioritized metabolites that were utilized by *C. difficile* based on the inferred *C_ij_* parameters in the CR model (fit to monoculture and community data) in DTL1 (205 unique conditions, 47 including *C. difficile*) and DTL2 (214 unique conditions, 37 including *C. difficile*). In DTL3 (285 unique conditions, 59 including *C. difficile*), we included metabolites underrepresented in previous DTL cycles to further refine our understanding of the metabolite design space. DTL4 (285 unique conditions, 44 including *C. difficile*) was partitioned into a group with metabolites used by *C. difficile* in communities (succinate, GlcNAc, trehalose, and mannitol) and a group with metabolites not used by *C. difficile* in communities as determined by the *C_ij_* parameter values in the CR model (sialic acid, sorbitol, GalNAc, galactose, mannose, and cellobiose). Bayesian experimental design was performed separately on each group. Detailed information about each DTL cycle is in **Table S5**.

To evaluate the improvement in model prediction performance across DTL cycles, we used a two-step cross-validation approach. When evaluating prediction performance of the model using data up to a particular DTL cycle, we first perform 20-fold cross-validation on all data collected up to that DTL cycle. In the second step, we train the model on all data up to that DTL cycle and use data from all subsequent DTL cycles as a test set. Combining the held-out predictions from cross-validation and the test predictions from subsequent DTL cycles provides model predictions for all samples in the entire data set. Model prediction performance is evaluated using the Pearson correlation coefficient and root mean square error (RMSE) between predicted and measured *C. difficile* abundances. By using predictions of all samples, we can evaluate how much incorporating data from subsequent DTL cycles improves model prediction performance in a way that ensures the performance metric is consistent across all comparisons. The DTL cycles were terminated at cycle 4 as RMSE had effectively plateaued, indicating diminishing returns from additional cycles **(Figure S4C)**. Over the four DTL cycles, the Pearson correlation coefficient of *C. difficile* abundance using the CR model moderately increased (0.48 in DTL0 to 0.52 in DTL4) **(Figure 2B and S4C)**. This implies that a significant fraction of the variance in the data was not fully captured by the CR model.

To provide deeper insights into how the Bayesian experimental design algorithm explores the metabolite-species design space across DTL cycles, we used multidimensional scaling (MDS) on community experiments inoculated with *C. difficile* to visualize the relationships between conditions. The conditions were encoded by binary vectors that capture presence or absence of species, metabolites and the phylogenetic relationships of species. In each DTL cycle, the algorithm selects experimental conditions on the edge of the design space **(Figure 2C)**. Across the characterized conditions, low *C. difficile* endpoint abundance was frequently observed **(Figure 2C)**, suggesting that there are potentially many mechanisms that can generate low *C. difficile* abundance.

A cluster of communities with high *C. difficile* abundance (cluster 4, mean *C. difficile* abundance 0.28 OD_600_, compared to 0.08 for cluster 0, 0.10 for cluster 1, 0.09 for cluster 2, and 0.12 for cluster 3) was identified using K-means clustering **(Figure S4D)**, with the number of clusters selected based on the sum of squared errors and average silhouette score **(Figure S4 E and F)**. *Eggerthella lenta* (EL), *Eubacterium hallii* (EH), and *Blautia hydrogenotrophica* (BH), spanning two different phyla, were more frequently in cluster 4 whereas *Bacteroides spp.* was infrequently observed in this cluster **(Figure S4G)**. Examining the nutrient environment of cluster 4 communities, we found that all metabolites were evenly represented, with the high amino acid condition present at a modestly higher proportion **(Figure S4H)**. Communities containing EL displayed significantly highest median *C. difficile* abundance across all DTL data compared to communities containing other species (p-value 2.76 ∗ 10^−6^) **(Figure S4I)**. This highlights robust positive impact of EL on *C. difficile* across various species-metabolite environments. By contrast, communities containing BH and EH did not display high median *C. difficile* abundances **(Figure S4I)**, suggesting that BH and EH have context-dependent enhancement of *C. difficile* growth in the high *C. difficile* cluster. In sum, our experimental design strategy uncovered a cluster with high *C. difficile* abundance, revealing key species that can positively impact *C. difficile* with varying degrees of robustness to species-metabolite perturbations.

### Model-guided biological insights into interactions that impact growth of C. difficile

A key advantage of mechanistic models over ML is the ability to examine parameters directly to gain deeper and causal insights into the interactions driving system behaviors. While CR models are typically parameterized from monoculture data^40,63^, resource utilization frequently shifts in a community context due to ecological interactions. Therefore, we used a CR model fit to data from all DTL cycles (CR-comm model) to generate biological insights into species–resource interactions. The *C_ij_* parameters fit to monoculture data represent the intrinsic capacity of a species to utilize a specific resource. By contrast, the inferred *C_ij_* parameters of the CR-comm model can be interpreted as the effective, context-averaged capacity of a given species to utilize a specific resource across diverse community contexts. The posterior distributions of species-specific death rates in CR-comm model included nonzero estimates for many species **(Figure S5A)**, indicating that this term in the model was useful for explaining trends in the species abundance data. In the CR-comm model, the inferred *C_ij_* for *C. difficile* were near zero for most resources and were enhanced for amino acids, succinate, and GlcNAc, indicating that *C. difficile* prefers these resources across all community contexts **(Figure 3A)**.

**Figure 3.**
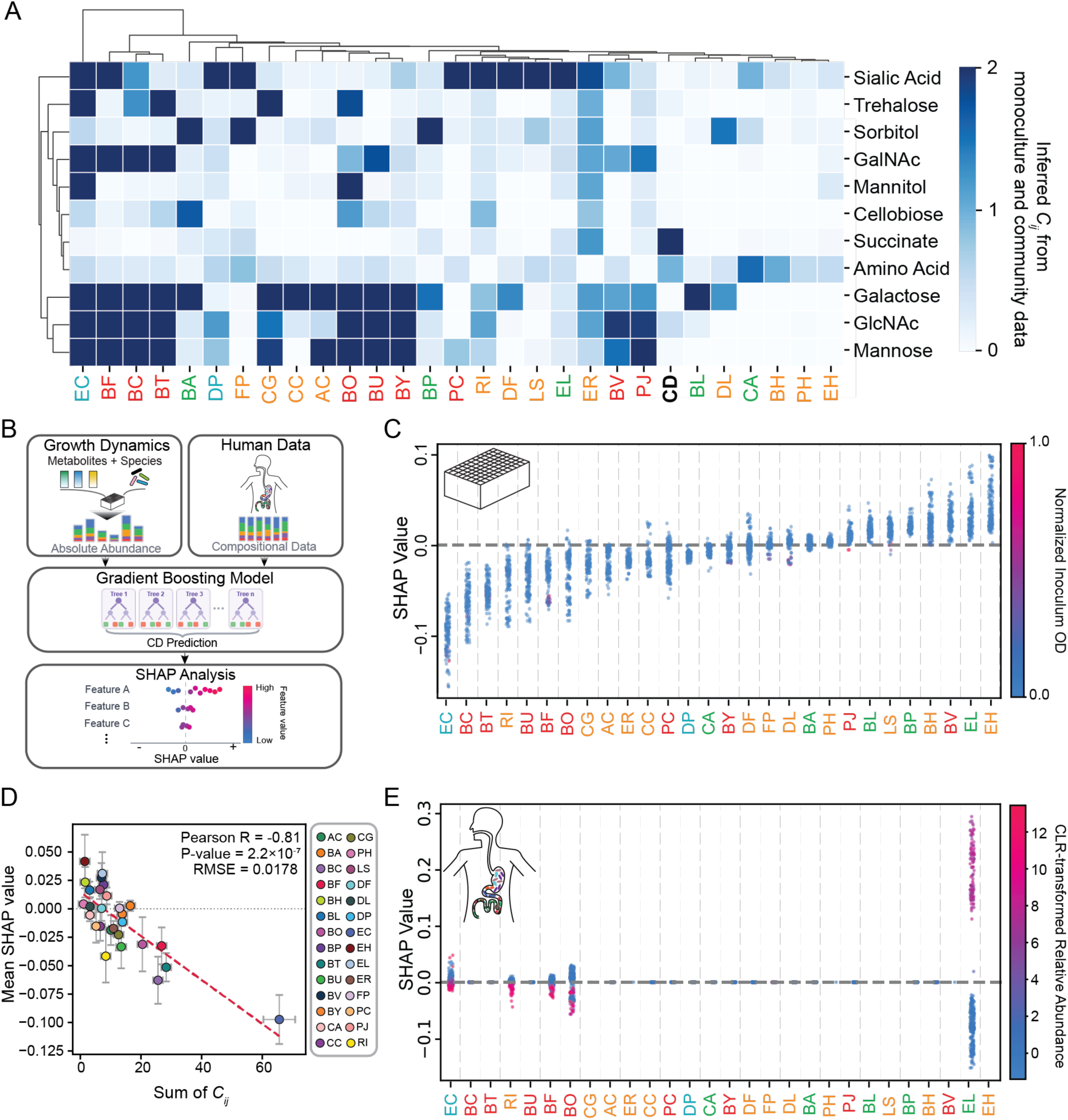
Model interpretation and biological insights from the CR and the GB model. **(A)** Clustered heatmap of inferred *C_ij_* parameters in the CR-comm model. The CR-comm model was trained on all DTL data, including both monoculture and community growth dynamics. Species colors represent phylum. The target species *C. difficile* (CD) is bolded and highlighted in black. **(B)** Schematic of GB models. GB models were independently trained on *in vitro* data from all DTL cycles and on human gut microbiome data^11,64^. For the *in vitro* model, inoculum species abundances and metabolite concentrations were used as input features to predict final *C. difficile* OD_600_. For the *in vivo* model, compositional data were used as input features to predict *C. difficile* presence or absence. Both models were coupled with SHAP analysis, an interpretable ML method, to quantify the contribution of each feature to *C. difficile* prediction. **(C and E)** Distribution of SHAP values from the *in vitro* model **(C)** and the *in vivo* model **(E)**. SHAP values were calculated across all conditions in which a given feature is present. Each data point represents the feature contribution in a single condition. Features are sorted by mean SHAP value from the *in vitro* model in descending order. Colors indicate normalized feature values: inoculum species OD_600_ for the *in vitro* model and centered log-ratio (CLR)-transformed relative abundance for the *in vivo* model. **(D)** Scatter plots of mean SHAP value versus summed *C_ij_* value across strains. Each dot represents an individual strain. The y-axis is the mean SHAP value from GB model for that strain, computed over the communities in which the strain is present (OD > 0). Error bars denote standard deviation of those SHAP values across communities. The x-axis is the sum of the strain’s positive consumption coefficients (*C_ij_* > 0) across all nutrients in the CR-comm model. Error bars denote ±1 standard deviation of the summed *C_ij_*, propagated from the posterior parameter uncertainty of the CR-comm model. The squared posterior standard deviations of the contributing coefficients (those with *C_ij_* > 0) were summed and the square root taken, i.e.[Inline]. The dashed red line is the linear least-squares fit. Colors indicate species identity.

The commensal species cluster into several groups including generalists (EC, BF, BC, and BT), specialists that utilize several mucin-derived sugars (CG, CC, AC, BO, BU, and BY), sialic acid (PC, RI, DF, LS, EL, and ER), or other variable patterns **(Figure 3A)**. The *C_ij_* parameter values for many species changed between the CR-mono model informed by monoculture data and CR-comm model fit to all monoculture and community data **(Figure S6A)**. Specifically, the *C. difficile*’s *C_ij_* parameters for all resources decreased in the CR-comm model, suggesting reduced utilization of all resources by *C. difficile* in community compared to monoculture. This implies that *C. difficile* may be partitioning its niche in communities to reduce the strength of inter-species competition. Therefore, metabolic niche overlap inferred from monoculture data does not accurately forecast *C. difficile* abundance in a community context. Consistent with this observation, the CR-mono model exhibited poor prediction performance of species abundance in communities and the CR-comm model displayed poor prediction performance on monoculture data **(Figure S5 B and C)**. However, informing the CR model with both monoculture and community data improved prediction performance of species abundance **(Figure S5 D and E)**, indicating that the effective, context-averaged *C_ij_* across monoculture and community data provides useful information to forecast community assembly.

The GB model, which maps initial species abundances and metabolite concentrations to *C. difficile* endpoint abundance, has the flexibility to capture a wide range of nonlinear and context-dependent interaction modalities that CR model does not capture. The context-dependent interactions represented in the GB model can be deciphered via explainable artificial intelligence methods like SHapley Additive exPlanations (SHAP)^50^. SHAP calculates the contribution of each feature to the model’s prediction.

Using all experimental data, we trained a GB model to predict the endpoint (24 h) *C. difficile* abundance from the initial species abundances and metabolite concentrations **(Figure 3B)**. The GB model achieves significantly higher prediction performance of *C. difficile* abundance (Pearson R = 0.77, **Figure S6B**) than the CR model (Pearson R = 0.52, **Figure 2B**) and a linear regression model with pairwise interactions (Pearson R = 0.72, **Figure S6C**), demonstrating that the GB model captures additional complexity not encompassed in the CR model. SHAP analysis generates a distribution of contributions from each input variable on the model’s prediction across all conditions in the dataset. Considering instances where the species or resource is present in the inoculum, a negative SHAP value indicates inhibition of *C. difficile* by the factor (species or resource), whereas a positive SHAP value indicates enhancement of *C. difficile* growth. Many species displayed strong inhibition of *C. difficile* across experimental conditions (e.g. EC, BC, BT, RI, BU, BF, and BO), while several species promoted *C. difficile* growth (e.g. PJ, BL, LS, BP, BH, BV, EL, and EH) **(Figure 3C)**. SHAP analysis of metabolite features revealed no individual carbohydrate as a strong predictor of *C. difficile* abundance **(Figure S6D)**. For amino acids, SHAP values were bimodal, reflecting the substantial increase in *C. difficile* abundance when amino acid concentrations were increased from 5 g/L to 10 g/L **(Figure S6D)**.

A comparison of the inferred interactions between the CR-comm model (i.e. *C_ij_*) and GB model (i.e. SHAP) can provide insight into the role of resource competition versus other interaction modalities in this system. We analyzed relationship between a given species’ SHAP value on the prediction of *C. difficile* endpoint abundance and this species’ *C_ij_* values across all metabolites in the CR models, representing its net resource competition capability across environmental contexts. The net resource consumption capability of each species displayed a strong negative correlation with mean SHAP values (CR-comm model, Pearson R = −0.81, P = 2.2×10⁻⁷, **Figure 3D**), whereas no informative relationship was observed for the CR-mono model (Pearson R = −0.17, P = 0.40, **Figure S6E**). The significant negative correlation for the CR-comm model indicates that the net consumption capability of each species across community contexts can explain a significant amount of variation in the SHAP values of the ML model. The negative correlation was robust to the removal of *E. coli*, which displayed the largest net resource consumption capability (CR-comm model, EC excluded, Pearson R = −0.73, P = 1.8×10⁻⁵, **Figure S6F**). By comparing the inferred mechanistic model parameters and explainable AI on the ML model, our results demonstrated that resource competition is a major driver of the trends in *C. difficile* abundance.

To assess whether the species-*C. difficile* interactions identified from *in vitro* data are also observed in the human gut, we trained a separate GB model on a longitudinal human study. A cohort of 79 individuals were treated with antibiotics and a low or high dose of a living bacterial therapeutic VE303 (defined synthetic consortium of eight bacterial strains) developed for prevention of rCDI, or placebo was administered to the individuals daily for 2 weeks **(Figure 3B, see Methods)**^11,64^. We used metagenomic sequencing-derived compositional data as input to predict the presence or absence of *C. difficile* on held-out test data, achieving a mean area under the ROC curve (AUC) of 0.88 based on 5-fold cross-validation, compared to about 0.5 for a null model with permutated labels **(Figure S6G)**. SHAP analysis was performed on the trained model using the detected bacterial species in the 29-member community as features **(Figure 3E)**. EC, RI, BF, and BO displayed negative SHAP values at high relative abundance, indicating negative association with *C. difficile*, whereas EL displayed a positive SHAP at high relative abundance, indicating positive association with *C. difficile* **(Figure 3E)**. These results are consistent with the sign of the SHAP values in the GB model trained on *in vitro* data **(Figure 3C)**, suggesting that key species–*C. difficile* interactions identified from *in vitro* experiments are also present in the human gut. In sum, these results suggest the potential translational relevance of our *in vitro* system to the human gut microbiome.

### Model-optimized communities inhibit C. difficile across different nutrient environments

Given the fluctuating and unpredictable nutrient conditions in the mammalian gut^33,34^, microbial communities that can consistently suppress *C. difficile* across distinct nutrient environments could be effective bacterial therapeutics. To evaluate the engineering capability of both models, we fit the CR-comm and GB models to all data and predicted the abundance of *C. difficile* at 24 h in all four-member communities in 22 nutrient environments **(Figure 4A)**. These conditions included the presence of all carbohydrates (10 added at 1 g/L, total 10 g/L) and leave-one-carbohydrate-out media (all but one added at 1 g/L, total 9 g/L) at two different amino acid levels (5 g/L and 10 g/L). To test the model predictions, we selected 15 communities with the lowest predicted maximum *C. difficile* abundance at 24 h across all simulated media conditions for experimental characterization in 22 media conditions (CR-opt and GB-opt for communities optimized using the CR-comm model and GB model, respectively). As a control, we selected 15 random communities (Random communities). Finally, we selected 15 communities predicted by both CR-comm and GB models to be among the 250 communities with the highest minimum *C. difficile* abundance across nutrient environments to identify diverse solutions (Low-Inhibition communities). The different model-designed sets of communities were used to evaluate the ability of the models to distinguish between inhibitory versus non-inhibitory *C. difficile* communities.

**Figure 4.**
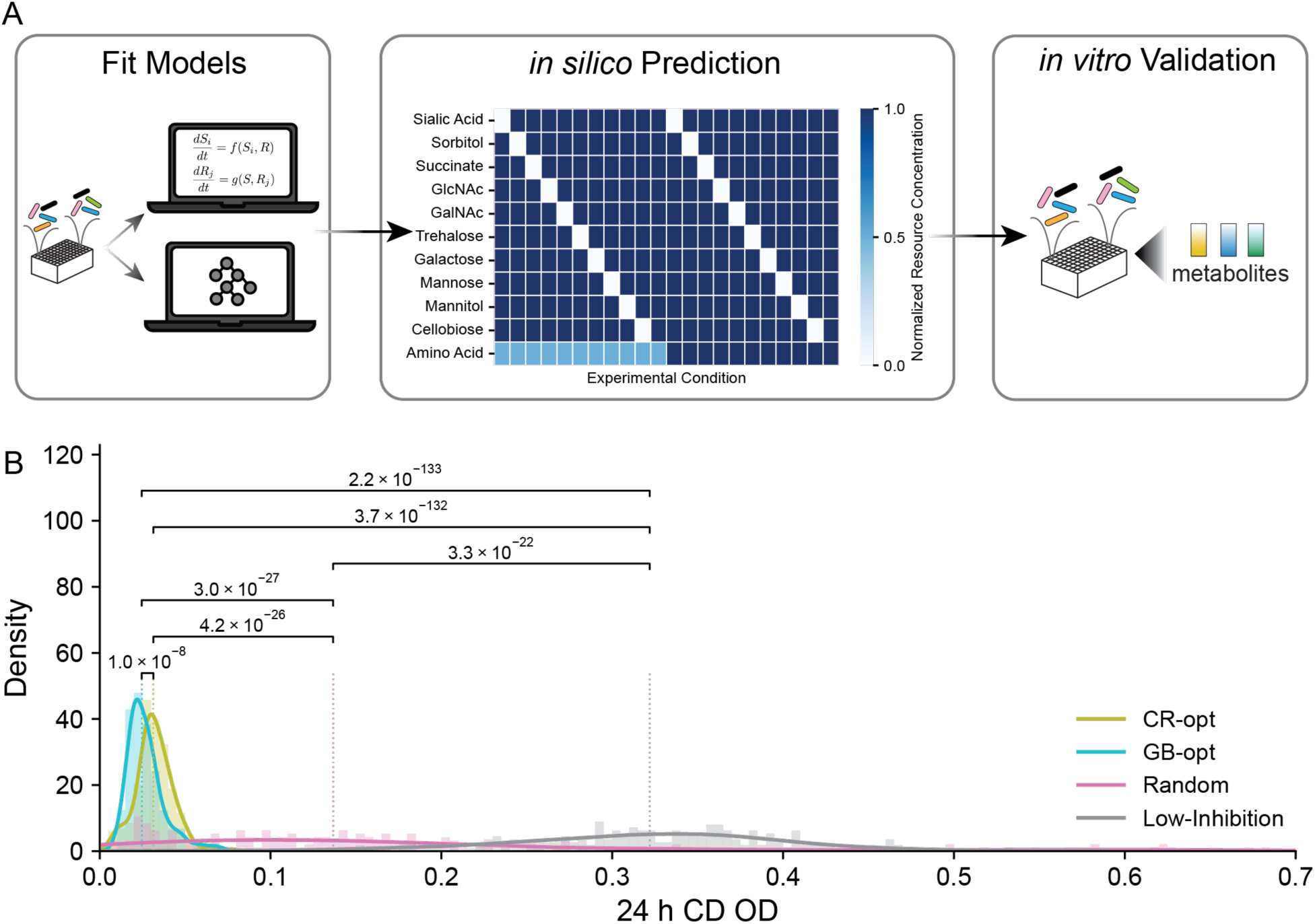
The *in vitro* validation of *C. difficile* inhibition by model-designed communities. (A) Schematic of model-designed community design and validation. All DTL experimental data were used to fit CR-comm model and GB model, separately. Each model was used to perform *in silico* prediction of 24 h *C. difficile* OD in all 4-member communities in 22 media conditions. The 22 media conditions (as illustrated by the heatmap) include full carbohydrates and leave-one-out carbohydrate conditions (1 g/L for each carbohydrate) at two different amino acid levels (5 g/L or 10 g/L). For higher *C. difficile* inhibition across metabolic environments, communities are ranked by maximum *C. difficile* OD across all *in silico* media conditions (as illustrated by the heatmap), with the lowest maximum *C. difficile* OD communities selected for validation. Fifteen communities with the lowest maximum *C. difficile* OD from each model were selected for validation (CR group and GB group). Fifteen communities with the highest maximum *C. difficile* OD from both models were also selected for validation (Low-inhibition group). Additionally, fifteen random communities were tested in validation (Random group). (B) Distribution of experimental 24 h *C. difficile* OD across all 22 media conditions. Each distribution includes a total of 330 unique experimental conditions from 15 communities in 22 media conditions. Statistical significance was assessed by unpaired t-tests.

Due to different inductive biases of the mechanistic and ML models, the predicted combinations of species that inhibit *C. difficile* can provide insights into different solutions for the design of anti-*C. difficile* communities^65^. EC is present in all model-optimized communities **(Figure S7 A and B)**, consistent with its large negative impact on *C. difficile* based on SHAP analysis using the GB model **(Figure 3C)** and the inferred high metabolic niche overlap with *C. difficile* in the CR-comm model **(Figure 3A)**. While EC was a universal feature, BC, BF, and BU frequently occur in CR-opt **(Figure S7B)**, whereas BO, BT, and CA are enriched in GB-opt **(Figure S7A)**. As expected, the Random communities consist of a uniform sampling of the species in the design space (**Figure S7C**). The Low-Inhibition communities are enriched with BL, BH, EH, and EL (**Figure S7D**). Consistent with this result, EH, BL, and BH are predicted to have low metabolic niche overlap with *C. difficile* in the CR-comm model **(Figure 3 A and D)** and positively impact *C. difficile* based on the SHAP analysis using the GB model (EL, EH and BH are within the top 4 highest values) (**Figure 3C**).

To evaluate the predictive capability of the models, we characterized all *in silico* predicted communities by assembling the communities in the same 22 media conditions used to simulate the model. We measured community composition based on 16S rRNA gene sequencing and total community biomass based on OD_600_. The mean absolute abundance of *C. difficile* was low in CR-opt and GB-opt communities compared to Random and Low-Inhibition communities (mean OD_600_ for *C. difficile* was 0.03 and 0.02 for CR-opt and GB-opt, respectively) across all media conditions **(Figure 4B)**, consistent with the model predictions **(Figure S8 A and B).** Notably, the GB-opt communities displayed significantly lower *C. difficile* abundance than CR-opt **(Figure 4B)**, implying that the GB model better captures key interactions influencing *C. difficile* growth. The Random communities displayed a broad range of *C. difficile* abundance and significantly higher mean (mean OD_600_ of 0.18) than CR-opt and GB-opt **(Figure 4B)**.

A key question is whether metabolic niche overlap alone is sufficient to design inhibitory communities. To this end, we computed the Jaccard index between *C. difficile* and each 4-member community using the inferred *C_ij_* parameters of the CR-mono model. Niche overlap based on monoculture growth exhibited no significant association with 24 h *C. difficile* growth OD _600_ in DTL0-4 (Pearson R = -0.032), suggesting that niche overlap alone does not account for the variation in *C. difficile* growth in community context **(Figure S8C)**. To further validate the inhibitory capacity of niche-overlap-selected consortia beyond this correlation analysis, we assembled additional 4-member communities with high or low niche-overlap that were not represented in our *in vitro* data (**Figure S8 D-F**). Among the high niche overlap communities, HO2 (BC, BT, BU, and EC; Jaccard index = 0.91) **(Figure S8E)**, which contains *E. coli*, partially inhibited *C. difficile* but was outperformed by GB1 (BF, BU, EC, and RI; Jaccard index = 0.91), an inhibitory community identified by optimizing *C. difficile* inhibition using the GB model **(Figure S8 H and I)**. By contrast, HO1 (BO, BY, LS, and PJ; Jaccard index = 0.82) failed to inhibit *C. difficile* **(Figure S8D)**, displaying *C. difficile* absolute abundance comparable to the GB model-designed non-inhibitory community 1 (EH, BH, BL, and EL; Jaccard index = 0.82) **(Figure S8 G and I)**. The low-overlap community LO (BH, CA, ER, and PC; Jaccard index = 0.18) also supported *C. difficile* growth to levels indistinguishable from non-inhibitory community 1, despite a nearly fivefold difference in their niche overlap (0.18 for LO vs. 0.82 for the non-inhibitory community 1) **(Figure S8 F, G, and I)**. These results demonstrate that communities selected based on monoculture-informed niche overlap displayed inconsistent effects on *C. difficile* growth, indicating this approach is unreliable for microbiome engineering.

Based on their initial metabolite concentration, species compositions, and phylogenetic relatedness, the GB-opt, CR-opt, Low-Inhibition, and Random conditions (validation set) were broadly distributed across the design space **(Figure S8J).** Therefore, there are multiple distinct solutions for *C. difficile* inhibition within the design space. To evaluate model prediction performance on these data, the trained models were used to predict *C. difficile* abundance in the validation dataset, which comprises all 60 experimentally characterized communities (CR-opt, GB-opt, Random, and Low-Inhibition) across 22 media conditions. Even though the distribution was multimodal due to the experimental design, the GB and CR-comm models displayed an informative relationship across all data in the validation set (Pearson correlation of 0.68 with p-value of 3.25 ∗ 10^−144^ for the CR-comm model, Pearson correlation of 0.66 with p-value of 4.40 ∗ 10^−132^ for the GB model) **(Figure S8 K and L)**. While the CR-comm and GB models displayed substantial differences in prediction performance, both models were identified communities that significantly inhibited *C. difficile* abundance *in vitro*. This implies that there is some flexibility in model prediction performance for optimizing desired functions^41^.

### Metabolic profiles and resource supplementation experiments identify key metabolites mediating C. difficile inhibition

To reveal the metabolic basis of *C. difficile* inhibition by model-designed communities, we selected four communities that displayed low *C. difficile* abundance across media conditions and exhibited differences in species presence/absence: CR1 (BO, BH, EC, and CA) and CR2 (BO, BT, EC, and FP) from CR-opt, and GB1 (BF, BU, EC, and RI) and GB2 (BC, CC, EC, and DF) from GB-opt **(Figure S9 A and B)**. To further validate the inhibitory performance of these communities, we compared *C. difficile* absolute abundance in these selected communities (GB1, GB2, CR1, and CR2) against non-inhibitory community 1 (BH, BL, EH, and EL) **(Figure S9 C and D)**.

Since the degree of inhibition of *C. difficile* by the model-designed communities could change over time, we performed a passaging experiment wherein every 24 hours an aliquot of the communities was transferred to fresh media (3 total passages) (**Figure S10 A-G**). The model-designed communities exhibited substantially lower *C. difficile* absolute abundance compared to CD-monocolonized or the non-inhibitory community 1, and inhibitory effects were maintained across 3 passages **(Figure S10 B-G)**. *C. difficile* absolute abundance estimated from the product of OD_600_ and relative abundance based on 16S sequencing was highly correlated to the *C. difficile* colony forming units (CFUs) **(Figure S10H)**, corroborating that the calculated absolute abundance metric accurately reflects viable cell counts.

*E. coli* was included in all four model-designed communities for *C. difficile* inhibition and was identified as an inhibitory species in the CR and GB models **(Figure 3 A, C, and D).** To examine the role of *E. coli*, we excluded *E. coli* from each community and quantified *C. difficile* abundance. *C. difficile* absolute abundance significantly increased in the absence of *E. coli* **(Figure S10I)**, demonstrating that *E. coli* plays a critical role in community-mediated inhibition.

To elucidate the metabolic mechanisms underlying *C. difficile* inhibition, we performed exo-metabolomic profiling on supernatant from monocultures, the four selected communities, and 16 leave-one-species-out sub-communities derived from these communities using UPLC-MS/MS **(Figure S11A) (see Methods)**. Peaks for GalNAc/GlcNAc, sorbitol/mannitol, galactose/mannose, and cellobiose/trehalose could not be resolved by uHPLC-MS and are therefore reported as metabolite pairs. In media with all 10 carbohydrates, *C. difficile* monoculture displayed significant utilization of GalNAc/GlcNAc, sialic acid, succinate, and sorbitol/mannitol **(Figure S11A)**. *C. difficile* monoculture also consumed a broad range of amino acids, including alanine, aspartic acid, cysteine, glycine, histidine, lysine, methionine, serine, threonine, and valine **(Figure S11A)**, consistent with the well-characterized Stickland amino acid fermentation pathways of *C. difficile*^66–71^. All model-designed communities exhibited significant utilization of GalNAc/GlcNAc, sialic acid, and sorbitol/mannitol **(Figure S11A)**. Since GB1, GB2, CR1, and CR2 communities successfully inhibited *C. difficile* **(Figure S9D)**, these metabolites may mediate resource competition with *C. difficile*. All communities lacking *E. coli* displayed similar metabolite profiles **(Figure S11A)**, indicating that *E. coli* had a major role in defining community-level metabolic profiles. Across GB1, GB2, CR1, and CR2, the metabolite profile of GB1 displayed the highest similarity *C. difficile* monoculture based on hierarchical clustering **(Figure S11A)**.

To test the contribution of resource competition on *C. difficile* inhibition in GB1, we performed sterilized supernatant experiments using GB1 in the presence and absence of nutrient supplementation. *C. difficile* growth was inhibited in the GB1 supernatant and carbohydrate and amino acid supplementation not only abolished the inhibitory effect but promoted *C. difficile* growth beyond fresh media levels **(Figure S11B)**. By contrast, nutrient supplementation in the supernatant of non-inhibitory community 1 did not significantly alter *C. difficile* growth **(Figure S11C**). In sum, our results demonstrate that the effects of the GB1 supernatant on *C. difficile* inhibition were mediated through resource competition.

To identify the contribution of individual species in GB1 on metabolite profiles, we performed time-series exo-metabolomic analysis of the GB1 community context with individual species removed. We defined a base community (BF, BU, and RI) lacking *E. coli* and evaluated changes in metabolite profiles with the addition of *E. coli* (EC+), *C. difficile* (CD+), or both species **(Figure 5 A and B)**. Comparing consumption profiles across conditions identified sorbitol/mannitol as the primary resource over which *E. coli* and *C. difficile* compete, as both species consumed sorbitol/mannitol when added to the base community **(Figure 5 C)**.

**Figure 5.**
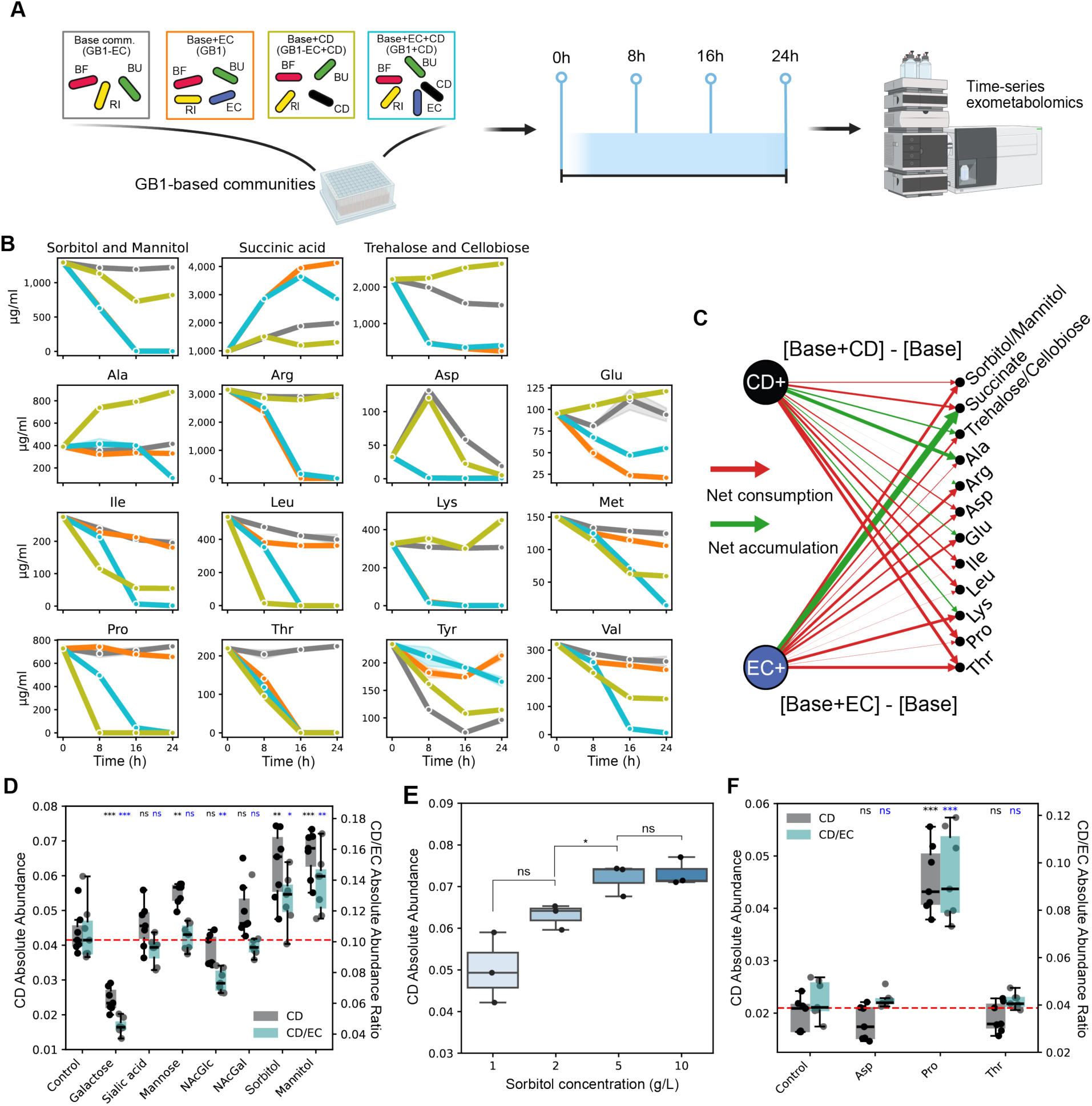
Leave-one-species-out exo-metabolomics analysis of the model-designed GB1 community. (A) Schematic of the leave-one-species-out time-series exo-metabolomics analysis. Starting from the base community lacking *E. coli* from GB1, *E. coli*, *C. difficile*, or both species were added individually or in combination. Culture supernatants were collected every 8 h and target metabolites were quantified by UPLC–MS/MS (n = 2). All species were grown in media containing 1 g/L each of 10 carbohydrates and 10 g/L amino acids. (B) Temporal profiles of metabolites whose concentrations changed by more than 50% at 24 h relative to initial (0 h) levels in the presence of *E. coli* or *C. difficile*. Colors indicate community composition: Base comm. (gray), Base comm. + *E. coli* (orange), Base comm. + *C. difficile* (olive), and Base comm. + *E. coli* + *C. difficile* (sky blue). (C) Bipartite network depicting species-specific metabolite consumption and synthesis by *E. coli* and *C. difficile* in the GB1 community context. Metabolites were defined as consumed or synthesized based on differences in concentration at 24 h between the Base comm. and communities with *E. coli* or *C. difficile*. Red and green edges denote consumption and synthesis, respectively; edge thickness represents mean normalized flux across conditions (n = 2). (D) Absolute abundance of *C. difficile* (left axis) and the *C. difficile*/*E. coli* absolute abundance ratio (right axis) under control conditions or upon 5× supplementation of carbon sources (n = 7). (E) *C. difficile* absolute abundance across increasing sorbitol concentrations (n = 3). (F) Absolute abundance of *C. difficile* (left axis) and the *C. difficile*/*E. coli* absolute abundance ratio (right axis) under control conditions or upon 5× supplementation of amino acids (n = 7). For all box plots, each dot represents an independent replicate; statistical significance was assessed by unpaired t-tests and is indicated as ns, not significant; *, P < 0.05; **, P < 0.01; ***, P < 0.001.

To assess whether the CR model captures the observed metabolite dynamics, we compared CR model predictions with time-series metabolomics measurements. This is a challenging task since the model was informed by species abundance and not metabolomics measurements and our CR model does not capture metabolite release, such as the net accumulation of succinate and trehalose/cellobiose observed in our experimental data **(Figure 5C)**. Nevertheless, the predicted metabolite concentrations followed similar temporal trends to the experimental measurements for most metabolites across all four community conditions **(Figure S12A)**. Succinate, trehalose/cellobiose, and total amino acids, which exhibited net accumulation were not accurately predicted by the CR model **(Figure S12B and C)**. After excluding these metabolites, the correlations for carbohydrates improved further **(Figure S12B)**, indicating that the CR model accurately captures resource consumption dynamics but not metabolite production, consistent with the model assumptions and structure.

Since sorbitol/mannitol were consumed in the presence of *E. coli* (EC+) and *C. difficile* (CD+) **(Figure 5C)**, we hypothesized that these metabolites mediate *E. coli*-driven *C. difficile* inhibition via resource competition. To test this hypothesis, we evaluated whether metabolite supplementation could relieve *C. difficile* inhibition *in vitro* in the context of the community. Galactose, sialic acid, mannose, GlcNAc, and GalNAc, were not consumed in the presence of *E. coli* or *C. difficile* and thus served as negative controls (supplemented with 5 g/L, five times the 1× control concentration (1 g/L) in fresh medium). Among these, mannose, sorbitol, and mannitol supplementation significantly increased *C. difficile* absolute abundance compared to the control **(Figure 5D, left y-axis)**. The ratio of *C. difficile* to *E. coli* absolute abundance (CD/EC ratio) was significantly increased only in the sorbitol-or mannitol-supplemented conditions compared to the control condition **(Figure 5D, right y-axis)**. Further, *C. difficile* absolute abundance increased proportionally and then saturated with increasing sorbitol concentrations **(Figure 5E)**. For amino acids, there was increased net consumption of Asp, Ile, Leu, Pro, and Thr in the CD+ and EC+ conditions **(Figure 5C)**. Supplementation of Pro (5X) reduced *E. coli*-mediated *C. difficile* inhibition, as evidenced by an increase in both *C. difficile* absolute abundance and the CD/EC abundance ratio **(Figure 5F)**. Collectively, exo-metabolomic profiling and resource supplementation experiments identified sorbitol, mannitol, and proline as potential key metabolites mediating *E. coli*-driven *C. difficile* inhibition through resource competition.

### Model-designed communities inhibit C. difficile colonization in a murine model

To evaluate consistency of model predictions of *C. difficile* abundance *in vitro* and in the mammalian gut, we performed *in vivo* experiments using germ-free mice. The *C. difficile* R20291 strain used for model training is a hypervirulent strain^72^ that can rapidly lead to the death of the mice, limiting our ability to assess longer-term changes in colonization ability. Therefore, we assessed the longer-term mammalian gut colonization dynamics using a reduced-virulence *C. difficile* strain MS008, which harbors *tcdA* and *tcdB* but lacks the binary toxin genes, *cdtA* and *cdtB*, associated with hypervirulence^73^. Despite the absence of binary toxin, the MS008 strain has been shown to occupy similar metabolic niches to hypervirulent strains^42^. To evaluate the robustness of *C. difficile* inhibition of the model-designed communities to variation in *C. difficile* genotype, we characterized the effects of inhibitory communities (GB1, GB2, CR1, and CR2) and non-inhibitory community 2 on the growth of *C. difficile* strains CD630^74^, R20291^75^ and clinical isolates MS001 and MS008^76^ using an *in vitro* 24 h co-culture experiment **(Figure S13A)**. All *C. difficile* strains exhibited significantly lower absolute OD when co-cultured with model-designed inhibitory communities compared to the non-inhibitory community 1 and *C. difficile*-only conditions, indicating that the *in vitro* inhibition strength of *C. difficile* did not vary substantially across isolates.

*P. hiranonis* and *L. scindens,* formerly classified within the Clostridium genus alongside *C. difficile*, have been consistently implicated in *C. difficile* inhibition^31,32,77^. Therefore, we considered a constrained optimization with the GB model for 4-member communities that maximize *C. difficile* inhibition across media conditions and contain *P. hiranonis* and *L. scindens*. Since *E. coli* was identified as a key contributor to *C. difficile* inhibition in our system, we designed communities with or without *E. coli*. With these constraints, the communities predicted to display the lowest *C. difficile* absolute abundance (24 h) across media conditions were BC, PH, LS, EC and BU, PH, LS, EC **(Figure 6A)**. These communities effectively inhibited *C. difficile* in media containing all 10 carbon sources (1 g/L each, total 10 g/L) and 10 g/L amino acids after 24 h of culture, displaying *C. difficile* absolute abundance moderately but significantly lower than GB1 **(Figure 6B)**. However, designed communities containing two closely related species but lacking *E. coli* (2Clo − EC) exhibited higher predicted *C. difficile* abundance compared to GB1 **(Figure 6A)**, consistent with its substantially weaker inhibition of *C. difficile in vitro* compared to GB1 **(Figure 6B)**. Similarly, the top-predicted four-member communities lacking *E. coli* (−EC) had weaker *C. difficile* inhibitory performance than GB1 in both model predictions and *in vitro* experiments **(Figure 6 A and B)**. Collectively, these results demonstrate that model-optimized communities constrained to include *P. hiranonis* and *L. scindens* can effectively inhibit *C. difficile*, consistent with previous studies^31,32,42^. These species inhibited *C. difficile* through non-bile acid dependent mechanisms^18,32^ since the culture medium did not contain primary bile acids **(Table S2)**. To identify candidate non-inhibitory communities for characterization in the mammalian gut, we used the model to identify communities that maximized *C. difficile* abundance across media conditions **(Figure 6A)**. Representative communities displayed significantly higher *C. difficile* absolute abundance compared to GB1 in *in vitro* experiments **(Figure 6B)**. Based on these results, we selected GB1, the constrained optimized community (BU, PH, LS, and EC), and non-inhibitory community 2 (BH, BV, EL, and PJ) for *in vivo* validation in germ-free mice.

**Figure 6.**
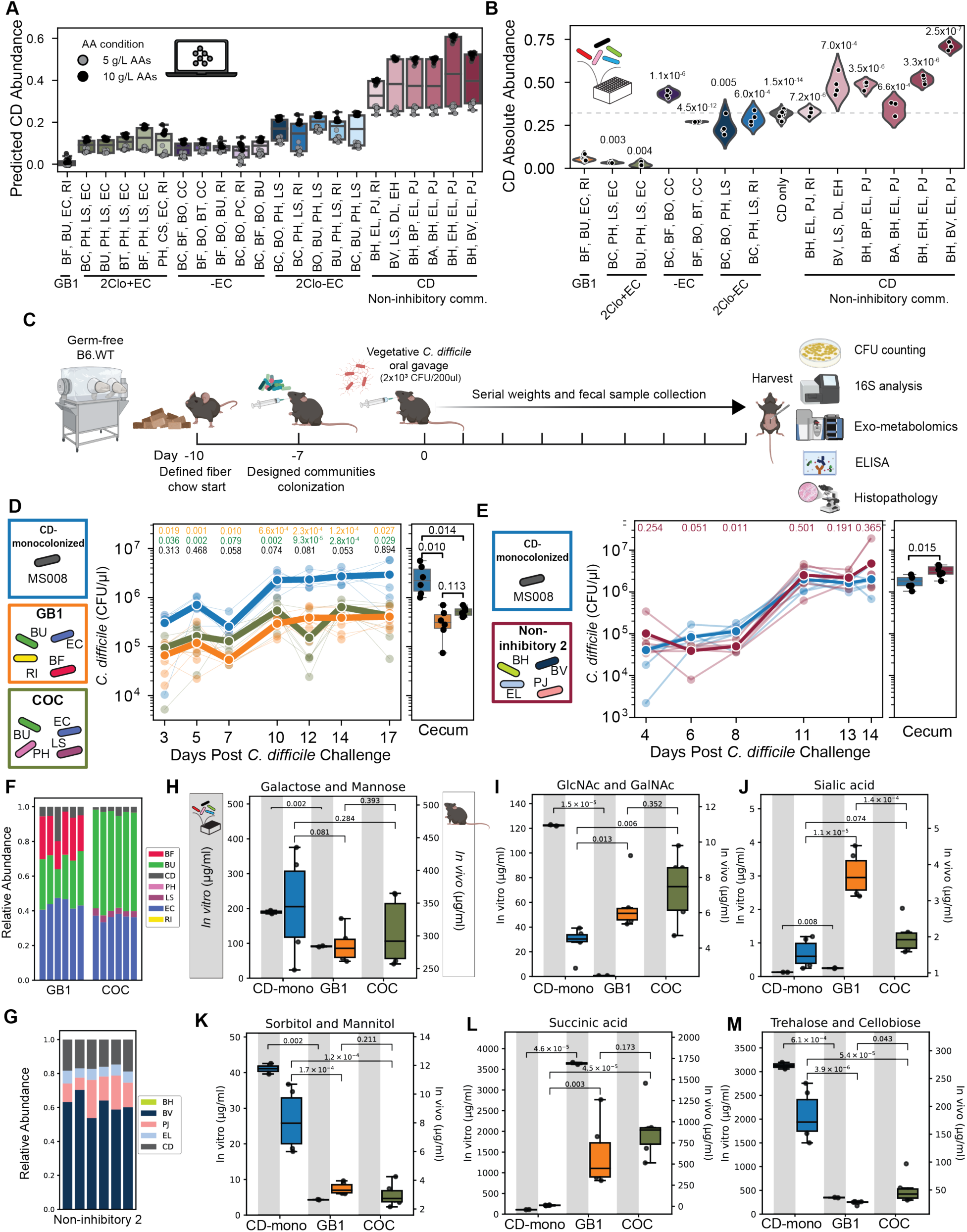
Model-designed communities inhibit *C. difficile* colonization in a murine model. (A) Boxplots of predicted 24 h *C. difficile* OD for designed communities across media conditions. Each dot represents the predicted *C. difficile* OD for a given community in a distinct media condition. Predictions were generated by the GB model across 22 media conditions: 11 carbohydrate conditions (all carbohydrates present and each of the 10 carbohydrates individually removed) at two amino acid concentrations (5 g/L and 10 g/L). 2Clo + EC, two *Clostridium* spp. with *E. coli*; 2Clo − EC, two *Clostridium* spp. without *E. coli*; −EC, without *E. coli*. Communities in the 2Clo + EC, 2Clo − EC, and −EC groups are sorted in descending order by the maximum predicted *C. difficile* OD across media conditions. *C. difficile* non-inhibitory communities are sorted in ascending order by the minimum predicted *C. difficile* OD across media conditions. (B) Boxplots of *C. difficile* absolute abundance in model-guided communities cultured in media containing 1 g/L of all 10 target carbohydrates with 10 g/L amino acids after 24 h. P-values from unpaired t-tests compared to GB1 are annotated above each boxplot (n = 8 for GB1, n = 4 for other communities). *C. difficile* MS008 strain was used. (C) Schematic of the murine experiment testing different model-designed communities. Mice were placed on a defined diet 3 days before oral gavage of human gut communities (GB1, COC, or non-inhibitory community 2). After 1 week of community colonization, 2000 CFU of *C. difficile* MS008 was introduced by oral gavage. Mice were weighed and fecal samples were collected every two to three days. After 2 weeks, mice were sacrificed and cecal contents were collected. Fecal and cecal samples were characterized via *C. difficile* colony forming units (CFU) counting and multiplexed 16S rRNA sequencing. Cecal samples were further characterized via *C. difficile* toxin ELISA assay and exo-metabolomics. Colon tissue samples were collected for histopathological analysis. (D and E) *C. difficile* CFU per μl of fecal and cecal samples. Panels D and E correspond to two independent murine experiments, each performed with its own CD-monocolonized control group: experiment 1 tested GB1 and COC (D), and experiment 2 tested the non-inhibitory community 2(E). For the line plots, colors represent experimental groups. Transparent lines represent biological replicates (each mouse) and solid lines represent the average across all biological replicates (n = 6 per group). Dashed line represents limit of detection. Statistical significance was compared between CD-monocolonized vs. GB1 (orange), CD-monocolonized vs. COC (dark green), and GB1 vs. COC (black) (D), and CD-monocolonized vs. non-inhibitory community 2 (E) by un-paired t-tests. P-values are annotated on each comparison. For the boxplots, *C. difficile* CFU per μl of cecal samples is shown. (F and G) Stacked barplots of species relative abundances in model-guided communities. (F) GB1 and COC. (G) Non-inhibitory community. Bar colors indicate species. (H-M) The barplots of exo-metabolomic analysis of cecal samples for Galactose/Mannose (H), GlcNac/GalNac (I), Sialic acid (J), Sorbitol/Mannitol (K), Succinate (L), and Trehalose/Cellobiose (M). For each condition, two bars are presented: the left bar represents *in vitro* results (gray background) and corresponds to the left y-axis, and the right bar represents *in vivo* results (white background) and corresponds to the right y-axis. *In vitro* samples were prepared from cultures in media containing 1 g/L of all 10 target carbohydrates with 10 g/L amino acids after 16 h (n = 2). *In vivo* samples were collected from cecal contents at the endpoint of the murine experiment (n = 6). Statistical significance was assessed by unpaired t-tests and p-values are annotated in the plots.

Germ-free female mice (8–16 weeks old) maintained on a standard grain-based chow diet were orally gavaged with each bacterial community or vehicle alone (PBS, no community) as a control. Germ-free mice were partitioned into four groups, including GB1, constrained optimized community (COC), non-inhibitory community 2, and no community group (*C. difficile* alone or CD-monocolonized) **(Figure 6C)**. The diet of the mice was switched three days before oral gavage of communities. One week after oral gavage with the communities (or no community), all groups of mice were orally gavaged with the same number of *C. difficile* vegetative cells. We used vegetative cells for consistency with our *in vitro* experimental design and computational model training as well as prior *in vivo* studies^69–71,78^. Fecal samples were then collected longitudinally, and cecal and colon tissues at the endpoint, to assess *C. difficile* abundance, community composition, toxin levels, metabolites, and histopathology **(Figure 6C; Methods)**.

The mice harboring GB1 or COC displayed significantly lower *C. difficile* CFU in across all fecal and cecal samples compared to the CD-monocolonized group, indicating that communities designed by the GB model effectively inhibited *C. difficile* colonization *in vivo* **(Figure 6D)**. By contrast, the *C. difficile* CFU in the non-inhibitory community 2 were similar to the CD-monocolonized in fecal samples and moderately and significantly higher in cecal contents than the CD-monocolonized group **(Figure 6E)**. Consistent with these results, *C. difficile* comprised on average 4.84 ± 2.93% and 3.54 ± 1.20% of the community in GB1 and COC groups, respectively, which were significantly lower than the 16.0 ± 1.76% observed in the non-inhibitory community 2 based on 16S sequencing **(Figure 6 F and G, S13 H and I)**. Collectively, these results demonstrate that model-designed communities can consistently inhibit *C. difficile* colonization in the mammalian gut over a two week period. Further, this inhibition property is specific to the model-optimized communities and not a general property of all 4-member communities as evidenced by the significantly higher abundance of *C. difficile* in the presence of the non-inhibitory community.

The normalized mice weights did not significantly change after introduction of *C. difficile* in any group **(Figure S13 C and D)**. In addition, ELISA quantification of total TcdAB concentration in fecal and cecal samples revealed no significant differences between any community group and the CD-monocolonized group **(Figure S13 E and F)**. Histopathological assessment of colon tissue samples showed no significant differences in inflammation total scores across groups **(Figure S13G)**. Collectively, these observations are consistent with the attenuated virulence profile of the MS008 strain^42,72,79^.

Because secondary bile acid production has been proposed as an alternative mechanism of *C. difficile* inhibition^31^, we additionally quantified bile acid concentrations in cecal contents by targeted exo-metabolomic profiling. Primary bile acids promote *C. difficile* spore germination and colonization, whereas the transformation of 7α-dehydroxylation into secondary bile acids has been shown to promote colonization resistance^26,31,80^. The transformation of 7α-dehydroxylation is restricted to a small number of subset of Clostridia that harbor the bile acid–inducible (*bai*) operon, such as *L. scindens*^81^. Primary bile acids were detected at significantly higher concentrations in mice colonized by GB1 and COC (mean 142.6 µM and 91.3 µM, respectively) than in the CD-monocolonized and non-inhibitory community 2 conditions (5.7 µM and 5.99 µM; **Figure S14A**), indicating that total primary bile acid abundance varied by colonized community. Secondary bile acids followed the same trend, reaching significantly higher concentrations in GB1 and COC (mean 0.10 µM and 0.10 µM) than in the CD-monocolonized and non-inhibitory community 2 conditions (0.01 µM and 0.02 µM; **Figure S14B**). However, these secondary bile acid concentrations were orders of magnitude below those required to inhibit *C. difficile* growth *in vitro* and *in vivo*^31,81,83^. Further, *L. scindens,* the only community member capable of secondary bile acid production through the *bai* operon^81^, was present at markedly low relative abundance even within COC **(Figure S13H)**. Together, these observations indicate that secondary bile acid production was not appreciable under our experimental conditions, suggesting that *C. difficile* inhibition by the model-designed communities was mediated predominantly by resource competition rather than by bile acid transformation.

To assess whether resource competition is consistent *in vitro* and *in vivo*, we performed targeted exo-metabolomic profiling of our design-space resources in cecal contents and compared them with *in vitro* measurements. Because absolute concentrations are not comparable across the two systems, we define a concordant consumption profile as one in which a resource shows the same direction of difference between a community-containing condition and the corresponding CD-monocolonized control in both *in vitro* and *in vivo* settings, whether depleted, unchanged, or accumulated. *In vitro* metabolomic profiling was performed for GB1 and the CD-monocolonized condition, whereas *in vivo* metabolomic profiling was performed for COC. *In vivo*, GB1 and COC showed similar carbohydrate profiles, differing only in sialic acid and trehalose/cellobiose **(Figure 6H–M)**. Relative to CD-monocolonized control, both communities showed lower sorbitol/mannitol and trehalose/cellobiose and higher GlcNAc/GalNAc, sialic acid, and succinate, whereas galactose/mannose was unchanged **(Figure 6H–M)**. For GB1, 7 of the 10 carbon sources showed concordant consumption profiles between *in vitro* and *in vivo* (galactose/mannose, sorbitol/mannitol, succinate, and trehalose/cellobiose). By contrast, mucin-derived sugars (GlcNAc/GalNAc and sialic acid) displayed inconsistent trends *in vitro* and *in vivo*. GB1 showed lower GlcNAc/GalNAc than CD-monoculture condition *in vitro* but higher *in vivo*, and sialic acid did not differ *in vitro* yet was higher in the communities *in vivo* **(Figure 6 I and J)**. This likely reflects the continuous, endogenous supply of mucin-derived sugars in the gut that is absent *in vitro*. Thus, for GB1, non-mucin-derived carbohydrates showed concordant consumption profiles across *in vitro* and *in vivo*, and COC mirrored the carbohydrate profile of the GB1 community *in vivo*.

For amino acids, GB1 and COC were nearly indistinguishable *in vivo* **(Figure S15)**. Relative to CD-monocolonized control, both communities enriched a broad set of amino acids (for example, alanine, glycine, branched-chain and aromatic amino acids, proline, serine, and threonine) and displayed lower concentrations of arginine, asparagine, and glutamine *in vivo* **(Figure S15)**. For GB1, only arginine, asparagine, glutamine, and proline showed concordant consumption profiles between *in vitro* and *in vivo*. The inconsistent trend in the majority of amino acids, unlike diet-limited carbohydrates^84,85^, may be because amino acids are also host-derived and synthesized endogenously by gut bacteria^31,37^, making their *in vivo* dynamics harder to predict from *in vitro* data.

Taken together, *in vivo* murine experiments demonstrated that the model-designed communities significantly inhibited *C. difficile* colonization in the mammalian gut, with non-mucin-derived carbohydrates showing concordant consumption profiles between *in vitro* and *in vivo,* whereas a subset of amino acids were consistent. By contrast, the other major mechanism known to inhibit *C. difficile*, secondary bile acids, remained orders of magnitude below inhibitory concentrations. These findings suggest that resource competition is the driving mechanism of the model-designed communities *in vitro* and *in vivo*.

## DISCUSSION

Recent clinical advancements underscore the potential of defined microbial consortia as therapeutics for *Clostridioides difficile* infection (CDI) that can address key disadvantages of FMT. VE303, a rationally selected consortium of eight *Clostridium* species, has demonstrated efficacy in a Phase 2 clinical trial for preventing recurrent CDI and is advancing through Phase 3^29,30^. Despite such progress, the mechanistic principles governing community-level inhibition of *C. difficile* remain incompletely understood, limiting our ability to rationally design and optimize living bacterial therapeutics. Existing approaches have largely relied on top-down strategies, such as screening reduced-complexity communities derived from healthy donor microbiota^38–40^. Rational selections of strains have leveraged phylogenetically related species that inhibit *C. difficile* through bile acid metabolism^41^ or non-toxigenic *C. difficile* strains for intraspecific competitive exclusion^41,42^. None of these approaches carefully considered the robustness of inhibition across environmental variability as a design feature. While each of these approaches has yielded promising results in specific contexts, we still do not fully understand which interaction mechanisms are most influential in inhibiting *C. difficile*. These knowledge gaps make it challenging to generalize inhibition strategies for other pathogens or microbiomes. A mechanistic understanding of how species and metabolites shape *C. difficile* growth and colonization across different nutrient landscapes would enable the rational design of synthetic microbiomes with broader therapeutic applicability.

Here, we developed a model-guided framework centered on resource competition, a prevalent modality in bacterial communities^84,85^. We evaluated a CR model that neglects cross-feeding model or a flexible GB model that can capture any interaction modality to predict community composition and design communities that robustly inhibit *C. difficile*. While the CR model displayed a moderate prediction performance (**Figure 2B)**, it was still successful in designing inhibitory communities *in vitro* **(Figure 4B)**. This implies that for certain target functions, maximizing prediction performance may not be critical for achieving desired microbiome engineering goals^85,86^.

Bayesian experimental design enabled efficient navigation of the vast species-metabolite design space by iteratively selecting maximally informative conditions across design-test-learn cycles **(Figure 1A, 2C, and S4)**^86,87^, prioritizing expected information gain over exhaustive sampling. This model-guided framework is particularly suited for microbiome engineering, where community-level behaviors emerge from context-dependent interactions that are not predictable from monoculture data alone, enabling principled navigation of combinatorial complexity with minimal experimentation. The GB model achieved better prediction performance for *C. difficile* abundance than the CR model, and GB-optimized communities displayed significantly lower *C. difficile* abundance **(Figure 4B and S6B)**. This is consistent with the flexibility of the machine learning model capturing interaction modalities beyond resource competition, such as cross-feeding and antimicrobial production^86^. This comparison should be interpreted with caution, however, as the two models differ in both inputs and outputs: the GB model predicts only endpoint *C. difficile* abundance, whereas the CR model predicts the temporal dynamics of all species, weighting each equally. SHAP analysis of the GB model revealed that species features exhibited strong and directionally consistent effects on *C. difficile* abundance **(Figure 3C)**, whereas individual resource concentrations showed no dominant effect on *C. difficile* outcome **(Figure S6D)**. This indicates that the composition of co-existing community members, rather than nutrient identity alone, is the primary determinant of *C. difficile* growth, consistent with resource competition being context-dependent based on constituent community members^87^.

*E. coli* emerged as a central contributor to *C. difficile* inhibition, consistently appearing in all model-optimized communities **(Figure 3 A, C and E; Figure 6 A and B; Figure S10I).** Among community members, *E. coli* exhibited the highest summed *C_ij_* across all metabolites in the CR-comm model **(Figure 3 A and D)** and exerted the greatest influence on the overall metabolic profile of the community **(Figure S11A)**. The *in vitro* exo-metabolomic profiling identified sorbitol, mannitol, and proline as candidate metabolites mediating this competitive interaction **(Figure 5 C-F)**. Sorbitol is particularly noteworthy, as it has been shown to promote both *C. difficile* expansion and virulence gene expression in the inflamed gut^88^. Given that *E. coli* depletes sorbitol in our *in vitro* system **(Figure 5 C-E)**, this raises the possibility that *E. coli*-mediated sorbitol depletion may not only attenuate pathogen growth but also virulence. Proline, a key substrate for Stickland fermentation^87^, alleviated *C. difficile* inhibition *in vitro* **(Figure 5F)**. Proline was one of the four amino acids (arginine, asparagine, glutamine, and proline) whose consumption profile observed *in vitro* was recapitulated *in vivo* **(Figure S15)**, whereas the other amino acids diverged between the two settings. We note that this *in vivo* comparison cannot isolate community-driven proline consumption, because *C. difficile* has been shown to consume proline and its abundance was substantially reduced in community-colonized mice than in the CD-monocolonized condition **(Figure 6D)**. The *in vivo* data therefore indicate that proline availability is altered in the presence of the designed communities, but the evidence that proline competition contributes to inhibition rests on the *in vitro* experiments and prior knowledge about its major role in mammalian gut colonization **(Figure 5F)**. Consistent with prior reports identifying proline as a critical resource for *C. difficile* colonization and fitness in the gut^88^, these findings support proline consumption as an additional competitive axis that may constrain *C. difficile*.

In the germ-free murine model, the GB1 community and COC which included *P. hiranonis* and *L. scindens*, commensal gut bacteria implicated in *C. difficile* inhibition^89^, significantly reduced *C. difficile* colonization **(Figure 6 D and F)**, whereas the model-predicted non-inhibitory community 2 failed to inhibit *C. difficile* colonization **(Figure 6 E and G)**. Exo-metabolomic profiling of cecal contents revealed concordant non-mucin-derived carbohydrate consumption profiles between the CD-monocolonized condition and GB1 across *in vitro* and *in vivo* settings **(Figure 6 H-M)**, suggesting that the resource competition observed *in vitro* is recapitulated *in vivo*^31,37,89^, and may contribute to *C. difficile* inhibition in the mammalian gut. Mucin-derived sugars showed divergent trends, likely reflecting continuous and endogenous supply from intestinal mucin absent *in vitro*^31,37,90^. We also examined secondary bile acid production, another potential modality of *C. difficile* inhibition^31,37,90^. Even in mice colonized by the inhibitory communities, secondary bile acids did not accumulate to sufficient concentrations to suppress *C. difficile* **(Figure S14B)**, indicating that resource competition, rather than secondary bile acids, was the major driver of inhibition by our designed communities. The inhibition of *C. difficile* by COC, which combines resource competition with potential bile acid-mediated inhibition through bile acids metabolism species^15,90^, suggests that integrating mechanistic resource competition-based design with other effective inhibitory mechanism may be a promising avenue for developing consortia that substantially inhibit *C. difficile* and exhibit robustness to environmental variability.

We limited our community design to communities of four species to reduce the design space and control for the previously demonstrated negative effect of species richness on *C. difficile* growth^15,91^. Relaxing this constraint in future work could likely discover defined communities with larger inhibitory potential. An open question, motivated by the diversity-invasibility relationship observed across microbial ecosystems^91^, is how many species are needed in a defined community to mirror the high efficiency of *C. difficile* inhibition by FMT. Further, our germ-free mouse experiments assessed only colonization resistance, with synthetic communities established prior to *C. difficile* challenge. Testing such treatment-mode inhibition, ideally within a naturally assembled gut microbiota rather than a germ-free background, represents an important direction for clinical translation. Finally, our design objective was *C. difficile* abundance, which is measurable at the throughput required for iterative design, rather than toxin production, which is more directly linked to pathogenesis. Incorporating toxin quantification into the active learning loop to select for communities that suppress toxin production would extend the framework to a more clinically relevant endpoint.

In conclusion, our model-guided design framework yielded synthetic communities that reduced *C. difficile* colonization significantly *in vitro* and *in vivo*, demonstrating that resource competition can be rationally engineered to suppress a clinically relevant pathogen. Combining such communities with dietary interventions that further shape the gut nutrient landscape in favor of inhibitory commensals offers another promising path toward pathogen suppression and elimination from the human gut. More broadly, we anticipate that this framework, complementary to existing top-down and empirically derived approaches, will extend to other pathogens and interaction modalities, accelerating the development of mechanism-informed microbiome-based medicines.

## Material and Methods

### Strain maintenance and culturing

All anaerobic culturing was carried out in anaerobic chambers with an atmosphere of 2 ± 0.5% H_2_, 15 ± 1% CO_2_ and balance N_2_. All prepared media and materials for anaerobic experiments were placed in the chamber at least overnight before use to equilibrate with the chamber atmosphere. The permanent stocks of each strain used in this paper **(Table S1)** were stored in 25% glycerol at −80 °C. Batches of single-use glycerol stocks were produced for each strain by first growing a culture from the permanent stock in ABB media **(Table S2)** to stationary phase, mixing the culture in an equal volume of 50% glycerol, and aliquoting 400 μL into Matrix Tubes (ThermoFisher) for storage at −80 °C. Quality control for each batch of single-use glycerol stocks included Illumina sequencing of 16S rDNA isolated from pellets of the aliquoted mixture to verify the identity of the organism. For each experiment, precultures of each species were prepared by thawing a single-use glycerol stocks (SUGs) and adding 100 μL of SUGs to 1 mL fresh ABB media **(Table S2)** for incubation at 37 °C for 24 or 48 h **(Table S1)**.

### Monoculture and community culturing dynamic growth quantification

Each species’ preculture was diluted to an OD_600_ of 0.01 (Tecan F200 Plate Reader) in a defined media^92^ designed in a previous study. Media pH is titrated to 7 using sodium hydroxide and hydrochloric acid, based on measurements obtained from a pH probe (Mettler-Toledo). Precultures were then aliquoted into three replicates of 1.5 mL each in a 96 Deep Well (96DW) plate and covered with a semi-permeable membrane (Diversified Biotech) and incubated at 37 °C without shaking. At each time point, samples were mixed by pipetting up and down 5 times before transferring 200 μL to 96 well microplate for measurement of OD_600_ by the plate reader (Tecan F200 Plate Reader). Cell pellets were spun down and saved at -80°C at each timepoint for species abundance quantification.

### Genomic DNA extraction from cell pellets

Genomic DNA was extracted from cell pellets using a modified version of the Qiagen DNeasy Blood and Tissue Kit protocol. First, pellets in 96DW plates were removed from −80 °C and thawed in a room temperature water bath. Each pellet was resuspended in 180 μL of enzymatic lysis buffer (20 mM Tris–HCl (Invitrogen), 2 mM sodium EDTA (Sigma-Aldrich), 1.2% Triton X-100 (Sigma-Aldrich), 20 mg/mL lysozyme from chicken egg white (Sigma-Aldrich)). Plates were then covered with a foil seal and incubated at 37 °C for 30 min with orbital shaking at 600 RPM. Then, 25 μL of 20 mg mL−1 Proteinase K (VWR) and 200 μL of Buffer AL (QIAGEN) were added to each sample before mixing with a pipette. Plates were then covered by a foil seal and incubated at 56 °C for 30 min with orbital shaking at 600 RPM. Next, 200 μL of 100% ethanol (Koptec) was added to each sample before mixing and samples were transferred to a nucleic acid binding (NAB) plate (Pall) on a vacuum manifold with a 96DW collection plate. Each well in the NAB plate was then washed once with 500 μL buffer AW1 (QIAGEN) and once with 500 μL of buffer AW2 (QIAGEN). Samples were then eluted into a clean 96DW plate from each well using 110 μL of buffer AE (QIAGEN) preheated to 56 °C. Genomic DNA samples were stored at −20 °C until further processing.

### Sequencing library preparation

Genomic DNA concentrations were measured using a SYBR Green fluorescence assay and then normalized to a concentration of 1 ng/μL by diluting in molecular grade water using either Tecan Evo Liquid Handling Robot or Hamilton STAR V. First, genomic DNA samples were removed from −20 °C and thawed in a room temperature water bath. Then, 1 μL of each sample was combined with 95 μL of SYBR Green (Invitrogen) diluted by a factor of 100 in TE buffer (Integrated DNA Technologies) in a black 384-well microplate. This process was repeated with two replicates of each DNA standard with concentrations of 0, 0.5, 1, 2, 4, 6, 8, and 10 ng/μL. Each sample was then measured for fluorescence with an excitation/emission of 485/535 nm using a Tecan Spark plate reader. Concentrations of each sample were calculated using the standard curve and a custom Python script was used to compute the dilution factors and write a worklist for either Tecan Evo Liquid Handling Robot or Hamilton STAR V to normalize each sample to 1 ng/μL in molecular grade water. Samples with DNA concentration <1 ng/μL were not diluted. Diluted genomic DNA samples were stored at −20 °C until further processing.

Amplicon libraries targeting the V3–V4 region of the 16S rRNA gene were prepared using one of two approaches. In the first approach, a single-step PCR was performed using custom dual-indexed primers **(Table S3)** arrayed in 96-well plates via an acoustic liquid handler (Labcyte Echo 550). In the second approach, a two-step PCR strategy was employed: the V3–V4 region was first amplified using custom primers **(Table S3)**, followed by a second PCR for unique dual indexing (Illumina UDI sets). For both approaches, PCR amplification was performed using Phusion High-Fidelity DNA Polymerase (Thermo Fisher) on normalized genomic DNA templates. PCR products were pooled, purified, and quantified prior to sequencing on an Illumina MiSeq platform using a 600-cycle kit to generate 2 × 300 paired-end reads.

### Bioinformatic analysis for quantification of species abundance

Sequencing data were demultiplexed using Basespace Sequencing Hub’s FastQ Generation program. Custom python scripts were used for further data processing^93^. Paired end reads were merged using FLASH (v1.2.11) after which reads without forward and reverse annealing regions were filtered out^94^. A reference database of the V3–V5 16S rRNA gene sequences was created using consensus sequences from next-generation sequencing data or Sanger sequencing data of monospecies cultures. Sequences were mapped to the reference database using the mothur (v1.48.3) command classify.seqs^95^. Relative abundance for each species within a sample was calculated as the read count mapped to each species divided by the total number of reads for each sample. Absolute abundance of each species was calculated by multiplying the relative abundance by the OD_600_ measurement for each sample. Samples were excluded from further analysis if >1% of the reads were assigned to a species not expected to be in the community (indicating contamination) or if they had < 1000 total reads and OD_600_ > 0.1 (indicating that there were insufficient reads for analysis and this was not due to lack of community growth).

### MDS embedding of community design space

Experimental conditions were represented as binary vectors capturing the presence/absence of species phylogeny and carbon sources. Phylogenetic information was augmented into each condition vector using parent nodes of the 16S phylogenetic tree, so that each of the 29 species was encoded as 57 phylogenetic-node indicators. Concatenating these with the 11 carbon-source indicators yielded a 68-dimensional binary design vector per condition. Multidimensional scaling was performed on the Euclidean distance matrix of these vectors and projected onto two dimensions using Scikit-Learn’s sklearn.manifold.MDS (SMACOF, random_state = 8).

### CR Model

CR models are ordinary differential equation models that can be used to model the dynamics microbial species growth on resources. Given a set of n species 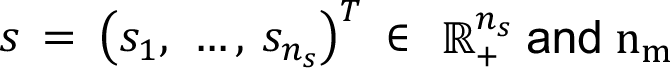 resources 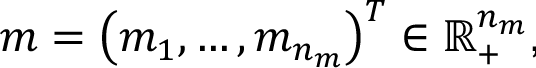 the proposed CR model is the following set of ordinary differential equations

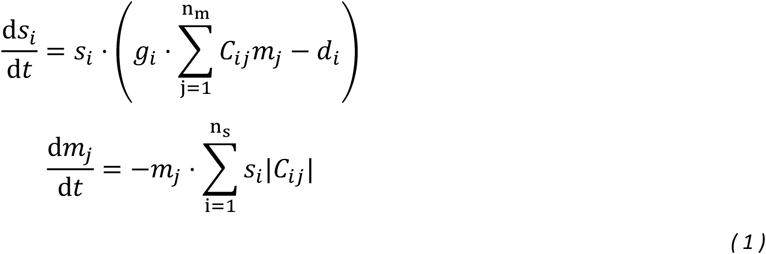

where the absolute value of *C_ij_* ∈ ℝ is the rate that species *i* consumes resource *j*, *g_i_* ∈ [0,1] is the species growth efficiency, and *d_i_* ∈ ℝ_+_ is the species death rate. We use the sigmoid function to constrain the species growth efficiency, where *g_i_* = 1/(1 + *e*^−*g*^^*^i^*), and we use a log transform to constrain the death rate to be positive where *d_i_* = 2*^d^*^^^*^i^*. If *C_ij_* < 0, then the consumption of metabolite *j* has a negative impact on the growth rate of species *i*. The set of unconstrained model parameters is denoted as θ = {*C_ij_*, *g^_i_*, *d^_i_*: *i* = 1, …, *n_s_*, *j* = 1, …, *n_m_*}. The absolute values of the fitted *C_ij_* parameters were used to characterize species resource consumption capacity in all downstream analyses, including resource utilization classification, metabolic niche overlap calculations, and clustered heatmap visualizations of conversion coefficients. We denote an experimental condition as *q*, which defines the species and resource initial conditions and measurement time after inoculation. An experimental measurement of species *i* abundance in experimental condition *q* is denoted as *y_i_*(*q*) and the corresponding model prediction is denoted as *f_i_*(*q*, θ).

### Defining the likelihood

We use the CR model to predict the expected value of an outcome from a particular experimental condition and characterize the noise as a Gaussian random variable as

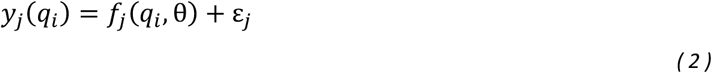

where 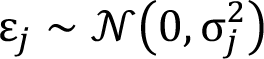 is a random variable that models measurement noise. Another way of expressing Eq. 2 is as a likelihood function,

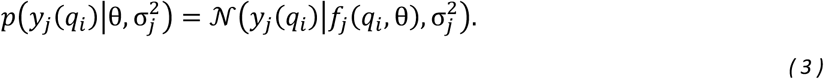

An experimental design is a set of experimental conditions, *q* = {*q*_1_, …, *q_n_*}, and data collected from an experimental design is denoted as D(*q*) = {*y*(*q*_1_), …, *y*(*q_n_*)}. Because measurements from each experimental condition are assumed to be drawn independently from Eq. 3, the likelihood of a set of measurements is

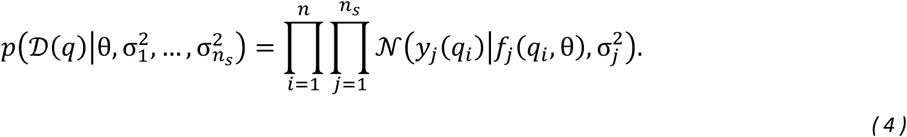

### Variational Inference and Expectation-Maximization for parameter inference and hyper-parameter optimization

The goal of variational inference is to optimize a parameter distribution so that it approximates the true parameter posterior distribution, *p*2θ|D(*q*)7. This is accomplished by optimizing the parameters, denoted as **φ**, of an approximate probability density function, *z*(θ|**φ**) ≈ *p*2θ|D(*q*)7. We will use an independent Gaussian prior distribution for each parameter, *p*(θ*_k_*|α*_k_*) = N(θ*_k_*|0,1/α*_k_*), where α*_k_* is the precision (inverse variance) of θ*_k_*.

The set of variables, 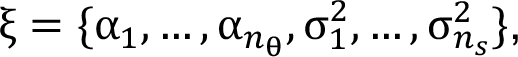 are considered hyper-parameters, because they influence parameter inference but are not explicitly a part of the model. These hyper-parameters control the degree of regularization, which is the amount of penalty placed on parameter values that deviate from the mean of the prior. Because the prior mean is zero in this case, increasing regularization promotes simplicity in the model in the form of zero-valued parameters. The objective function that takes into account model hyper-parameters, ξ, is the log marginal likelihood given by

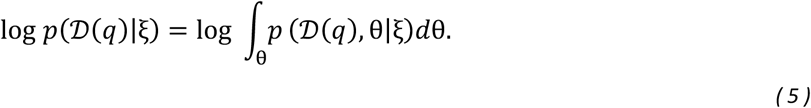

For any choice of *z*(θ|**φ**), the log marginal likelihood can be decomposed into two functions,

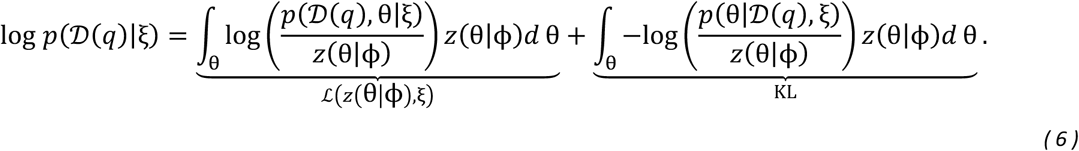

where KL is the Kullback-Leibler divergence between the approximating distribution *z*(θ|**φ**) and the true posterior parameter distribution *p*(θ|D(*q*), ξ). Because the KL divergence is strictly non-negative, the function ℒ(*z*(θ|**φ**), ξ) is a lower bound on the log marginal likelihood since ℒ(*z*(θ|**φ**), ξ) ≤ log *p*(D(*q*)|ξ). This lower bound is referred to as the evidence lower bound (ELBO). Optimizing the ELBO with respect to the parameters of the approximate posterior distribution, **φ**, is called variational inference. The Expectation-Maximization algorithm iterates between an expectation step, which involves variational inference to maximize the ELBO with respect to **φ** keeping ξ fixed, followed by a maximization step, in which the ELBO is maximized with respect to ξ keeping **φ** fixed.

### Expectation step for independent Gaussian parameter posterior

To enable more scalable Bayesian inference, we can use a simpler distribution to approximate the true parameter posterior where parameters are assumed to be independent, *z*(θ|**φ**) = 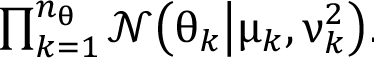 Evaluating the ELBO in this case gives,

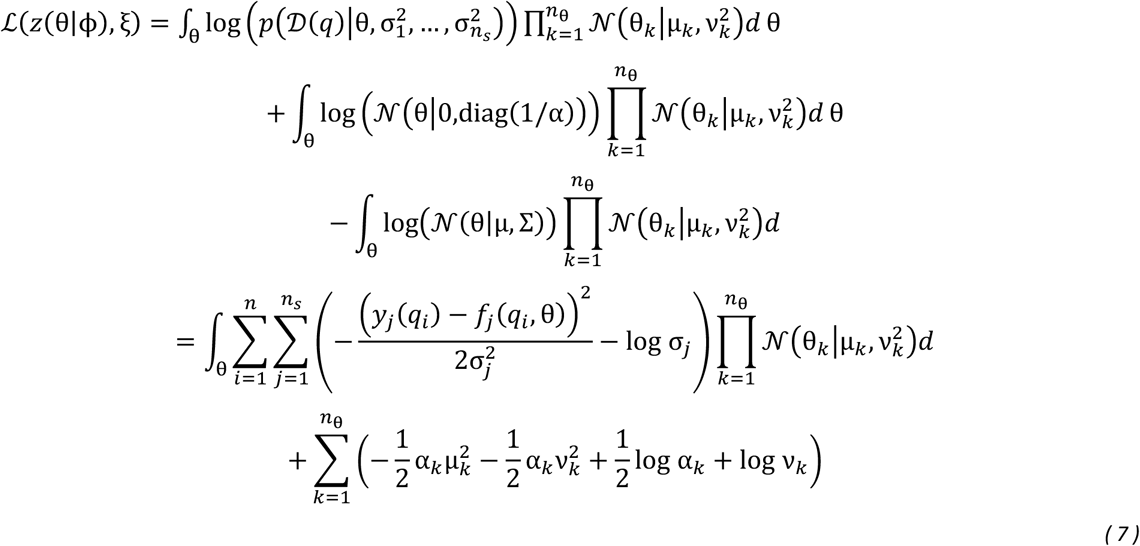

Linearizing the model with respect to parameters about the point θ = µ gives,

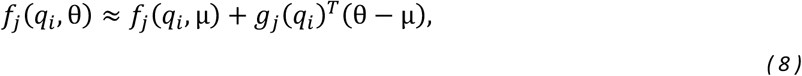

where we define *g*_j_(*q_i_*) = ∇_θ_*f*_j_(*q_i_*, θ). Using the linearized model to evaluate the expectation in the ELBO gives,

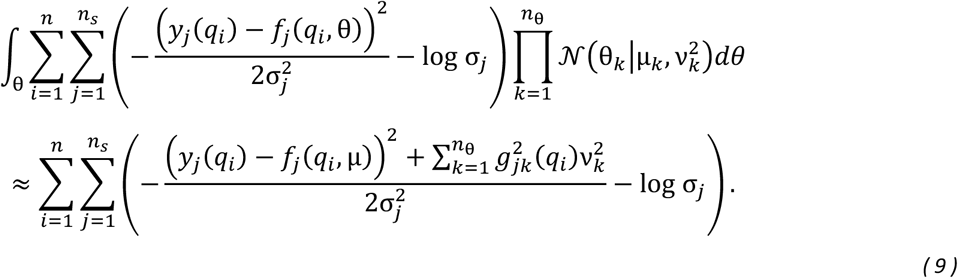

Putting all of the terms together gives the approximate expression for the ELBO,

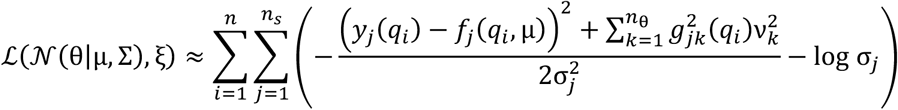

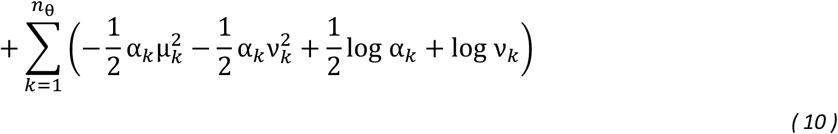

Maximizing this expression for the ELBO with respect to $\mu$ gives the same estimate as before,

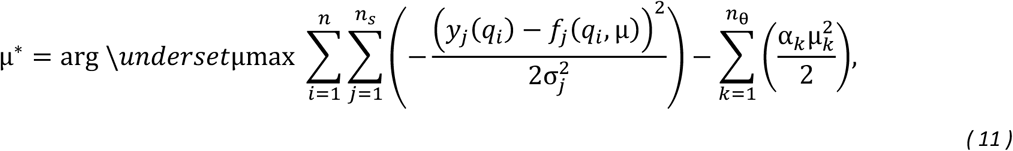

and using the estimate for µ^∗^, maximizing the ELBO with respect to ν^2^ gives,

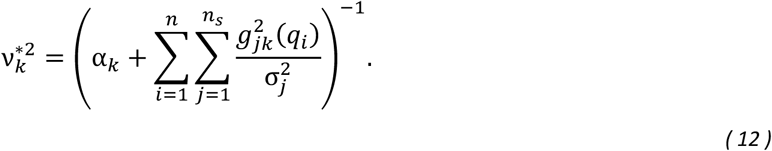

Optimization of µ is performed using the Newton-CG method from the Scipy minimize function^96^.

### Maximization step for independent Gaussian parameter posterior

Maximizing Eq. 10 evaluated using the optimized variational parameters with respect to each α*_k_* and each σ^2^ now gives

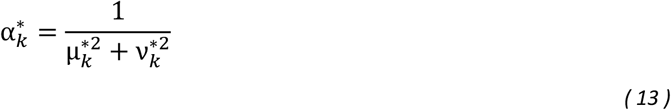

and

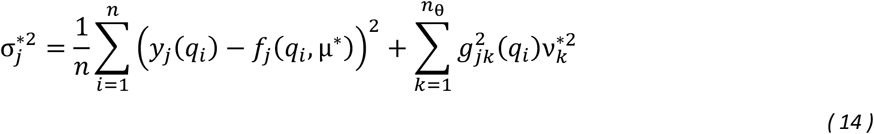

### Wald test for parameter significance

For a parameter θ*_i_*, we use a two-tailed Wald test to test whether the parameter is significantly constrained to be non-zero by the experimental data. Intuitively, the Wald test compares the parameter mean to its standard deviation in order to evaluate whether the peak of the posterior parameter distribution is significantly higher or lower than zero compared to the width of the distribution. We denote the null hypothesis as, *H*_0_: θ*_i_* = 0. The Wald test of size α = .05 for parameter *i* rejects *H*_0_ when |*W_i_*| > *z***_α_**_/2_, where

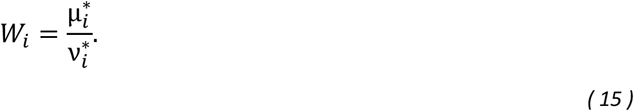

The p-value for an observed Wald statistic, *W_i_*, is given by *p* = 2Φ(−|*W_i_*|), which is the probability of the Wald statistic being at least as large as the observed value under the null hypothesis^97^.

### Estimation of posterior predictive distribution

Given a posterior parameter distribution, the posterior predictive distribution is found by marginalizing model predictions with respect to parameters,

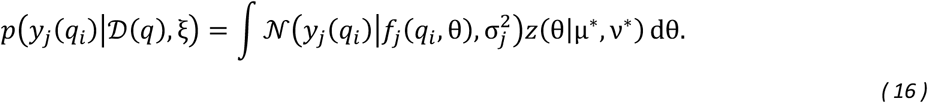

Linearizing the model prediction with respect to the model parameters around µ^∗^ allows for the following approximation of the integral in Eq. 16.

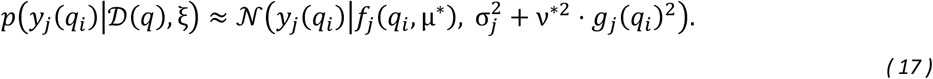

This approximation is referred to as the linear-Gaussian approximation^33,98^.

### Bayesian experimental design

The goal of Bayesian experimental design is to select an experimental design, denoted as *q* = {*q*_1_, …, *q_n_*} that results in data, D(*q*) = {*y*(*q*_1_), …, *y*(*q_n_*)}, that is expected to minimize the spread of the parameter distribution according to Bayes’ formula,

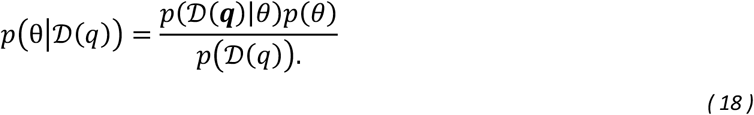

The variable *q_i_* is introduced to denote a particular experimental condition, which could for example be the specification of species to inoculate and resources to include in the media. It is important to note that the prior *p*(θ) could be the posterior determined from performing inference on previous data. In order to select an optimal experimental design, we define a utility function, *U*(*q*), as the expected information gain (EIG) that would result from updating the current model with a new dataset D(*q*). The EIG is the expected Kullback-Leibler divergence between the parameter posterior and the parameter prior distributions, which has an analytical expression in the case of the linear-Gaussian model

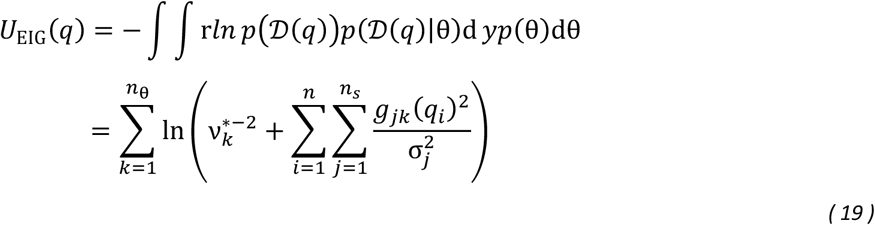

where 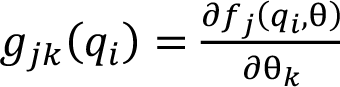 is the gradient of the model prediction of experimental condition *q* with respect to model parameters.

Given a set of all possible experimental conditions, denoted as Q, we use a Greedy search algorithm to determine the subset of *n* experimental conditions that maximize Eq. 19, where the first selected experimental condition is given by,

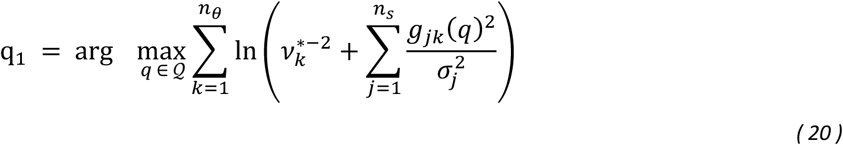

and subsequent experimental conditions are selected according to

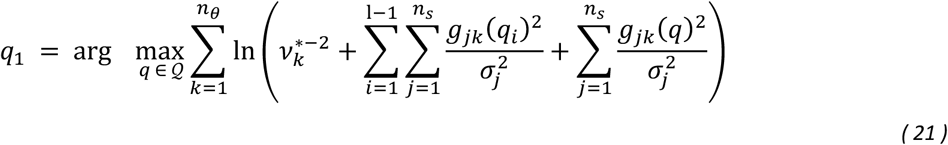

until l = *n*, where *n* is the total number of conditions that are experimentally practical to collect.

### Modified linear regression model

A modified linear regression model was used to assess the effect from inoculum species and resources on *C. difficile* abundance at 24 h. This model assumes linear influence of the covariates (inoculum species OD and resource concentration) and interactions among the covariates to the variable of interest (*C. difficile* OD at 24 h)^99^. The model formulation is as follows:

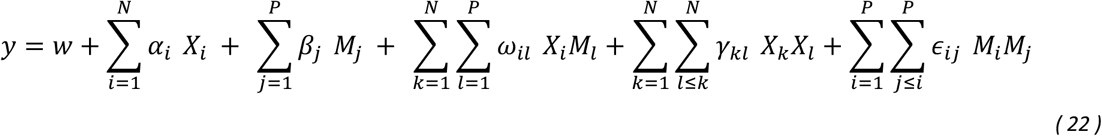

Where *X* ∈ ℝ*^N^*^×*S*^represents inoculum species OD and *M* ∈ ℝ*^P^*^×*S*^represents inoculum resource concentration. In this model we aim to predict *y* ∈ ℝ*^S^*, the OD of *C. difficile* at 24 h. Each parameter will provide insights into the connection of each covariate towards the growth of *C. difficile*. Specifically, *α_i_* signifies the influence of species *i*, and *β*_j_ provides insights into the influence of the nutrient *j*, *ω_il_* quantifies the effect of the co-occurrence of organism *i* and nutrient *l*, *γ_kl_* quantifies the effect of the co-occurrence of organisms *k* and *l,* and *ε_i_*_j_ the effect of the co-occurrence of nutrient *i* and *j.* This model was developed using the scikit-learn framework^49,100^. Model prediction performance was assessed by 10-fold cross validation performed over the whole dataset, splitting the dataset using the ShuffleSplit method of sklearn.

### Gradient boosting models and SHAP analysis

Gradient boosting is an ensemble method that sequentially constructs regression or classification trees, where each new tree corrects the residuals of the preceding ensemble^100^. We used two gradient boosting implementations depending on the dataset: LightGBM for the *in vitro* data and XGBoost for the *in vivo* data. For the *in vitro* analysis, the LightGBM model was trained to predict the endpoint (24 h) *C. difficile* absolute abundance, defined as the product of total OD_600_ and *C. difficile* relative abundance, from the initial OD of each constituent species and metabolite concentrations in the inoculum. For the *in vivo* analysis, XGBoost was applied to metagenomic data from the Vedanta dataset^101^, which contains longitudinal microbiome profiles from *C. difficile*-infected patients who received antibiotics followed by either a synthetic fecal microbiota transplant or placebo over a 168-day observation period. Microbiome data were filtered to retain species present in >1% of samples and central log-ratio transformed. A binary outcome variable, CDIFF_PRESENCE, was defined based on the detection of *C. difficile* in each sample. The hyperparameters of the models were tuned via the Optuna framework^101^. For insight into species and resource contribution to *C. difficile* growth, we adopted a ML interpretation method named the SHapley Additive exPlanation (SHAP) method^102^. This method gives us the faculty to assign negative and positive values to covariates that in a certain experiment contribute to the movement of the prediction against the mean values, therefore we can infer which microorganisms or nutrients have negative or positive contribution on the *C. difficile* outcomes. Calculations of SHAP values are performed through the SHAP Python package.

### Untargeted metabolomics by uHPLC-MS

Metabolite profiling of community culture supernatants was performed using an ultrahigh-pressure liquid chromatography–mass spectrometry (uHPLC-MS) system consisting of a Thermo Scientific Vanquish uHPLC coupled to a heated electrospray ionization source and a Q Exactive Orbitrap mass spectrometer (Thermo Scientific). The ion source was operated in negative polarity with the following settings: auxiliary gas flow rate of 10, sheath gas flow rate of 30, sweep gas flow rate of 1, spray voltage of 2.5 kV, capillary temperature of 320°C, and S-lens RF level of 50. Nitrogen was used as the nebulizing gas. Chromatographic separation was achieved on a Waters Acquity UPLC BEH C18 column (2.1 × 100 mm, 1.7 μm particle size). Mobile phase A consisted of 97% water and 3% methanol with 10 mM tributylamine adjusted to pH 8.2 with glacial acetic acid, and mobile phase B was 100% methanol. The gradient was as follows: 5% B from 0–2.5 min, linear increase to 95% B from 2.5–17 min, hold at 95% B for 2.5 min, return to 5% B by 20 min for re-equilibration, at a constant flow rate of 200 μL/min. The autosampler temperature was 4°C, injection volume was 10 μL, and column temperature was 30°C. Full-scan MS1 data were acquired from 70 to 1,000 m/z at a resolution of 70,000. Metabolite quantification was achieved using standard curves generated from six-point serial dilutions of authentic standards (0.01–10 mM; Sigma-Aldrich) dissolved in HPLC-grade water and stored at −20°C. Samples were diluted 1:100 or 1:1,000 so that target analyte signals fell within the linear range of the standard curves. The following pairs of isomeric metabolites could not be chromatographically resolved and are reported as combined concentrations: sorbitol/mannitol, galactose/mannose, N-acetylglucosamine/N-acetylgalactosamine, and trehalose/cellobiose. Resource utilization was calculated as:

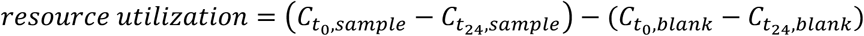

where C represents the measured absolute metabolite concentration in either sample or blank at 0 h (t₀) or 24 h (t₂₄). Normalized fold change was calculated as:

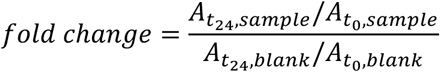

where A represents the area under the curve in either sample or blank at 0 h (t₀) or 24 h (t₂₄).

### Targeted metabolomics by UPLC-MS/MS

Targeted quantification of carbon source compounds, amino acids, and bile acids was performed at the Duke Proteomics and Metabolomics Core Facility using ultra-performance liquid chromatography–tandem mass spectrometry (UPLC-MS/MS) with multiple reaction monitoring (MRM). Carbon source and amino acid assays were run on a Sciex QTrap 6500+ mass spectrometer coupled to a Waters Acquity I-class Plus UPLC system, with calibration curves prepared in accordance with the FDA Guidance for Bioanalytical Method Validation101; data were acquired using Analyst 1.7.3 (Sciex) and processed in Skyline (v23.1)102. Across all assays, calibration standards spanning the assay-specific range, quality control samples, and study pool quality control (SPQC) samples were included to monitor accuracy and analytical consistency. For carbon source analysis, leave-one-species-out community supernatants were thawed on ice, vortexed, and centrifuged (10,000 × g, 5 min, 4°C). Supernatant aliquots (20 μL) were diluted with 40 μL water in 96-well plates, and calibration standards and QC samples were prepared by serial dilution in blank DM38 medium. Chromatographic separation used a Waters Acquity CSH Phenyl-Hexyl column (2.1 × 100 mm, 1.7 μm) with 5 mM ammonium formate in water (mobile phase A) and acetonitrile (mobile phase B) at 40°C. Spectra were acquired under negative electrospray ionization (ESI−) at a spray voltage of −4,500 V and source temperature of 450°C.

For amino acid analysis, supernatants stored at −80°C were thawed, vortexed, and centrifuged (15,000 × g, 2 min, 10°C). Aliquots (5 μL) were derivatized with benzoyl chloride (2% in acetonitrile) in the presence of 100 mM sodium carbonate following established benzoyl chloride labeling protocols103, and 13C6-labeled amino acid internal standards were spiked into each sample. Separation used a Waters Acquity UPLC BEH C18 column (2.1 × 50 mm, 1.7 μm) with 0.1% formic acid and 5 mM ammonium formate in water (mobile phase A) and acetonitrile (mobile phase B) at 40°C. Spectra were acquired under positive ESI at 3,200 V and source temperature of 500°C. Twenty amino acids were quantified using compound-specific MRM transitions with corresponding 13C6-labeled internal standards.

For bile acid analysis, bile acids in murine cecal contents from the gnotobiotic experiment were quantified using the Biocrates Bile Acids targeted assay. Cecal contents were diluted in isopropanol at 20 µL per mg of material and homogenized by bead beating, and the cleared supernatant was used for analysis. Sample registration, data acquisition, and quantification were managed in Biocrates MetIDQ software, with absolute concentrations calculated from calibration curves of authentic standards. Three levels of quality control samples (QC1–QC3) were injected in triplicate on every plate, and analyte-specific limits of detection were determined per plate.

### Gnotobiotic mouse experiments

All germ-free mouse experiments were performed following protocols approved by the The Duke Gnotobiotic Core, managed by the Division of Laboratory Animal Resources (DLAR). The defined diet used for all experiment was Envigo 2018S, containing 18.4% protein, 6.0% fat, 44.2% carbohydrate, 3.8% crude fiber, 14.7% neutral detergent fiber, and 5.5% ash **(Table S4)**. All strains were grown at 37 °C anaerobically in ABB media (Oxoid) for 24 or 48 h **(Table S1)**. All strains for oral gavage were mixed in equal proportions based on OD_600_ and transferred on ice prior to oral gavage. We used 8-16 weeks old C57BL/6 gnotobiotic female mice (wild-type) fed the defined diets 3 days prior to oral gavage. The mice from the same group (6 mice) were housed in the same biocontainment cages (Allentown Inc.) for the duration of the experiment. Mice were maintained on autoclaved water. One week after community gavage, 2000 CFU of vegetative *C. difficile* MS008 cells were orally gavaged into all mice in all groups. Mice were weighed and fecal samples were collected every 2-3 days after oral gavage for NGS sequencing and *C. difficile* CFU counting on TCCFA selective media^103^. About two weeks after *C. difficile* challenge, mice were euthanized, and the cecal contents were collected for NGS sequencing, *C. difficile* CFU counting, and toxin ELISA assay. For histopathology analysis, the entire colon from each animal was collected in 10% neutral buffered formalin (Sigma-Aldrich), fixed for 48 hours, processed, embedded, and sectioned routinely, and stained with hematoxylin & eosin. Sections were scored by a board-certified veterinary pathology according to published recommendations for histomorphological scoring of colitis in murine models^104^.

### Genomic DNA extraction from fecal and cecal samples

The DNA extraction for fecal and cecal samples was performed as described previously with some modifications^104^. Fecal samples (∼50 mg) were transferred into solvent-resistant screw-cap tubes (Sarstedt Inc) with 500 μL 0.1 mm zirconia/silica beads (BioSpec Products) and one 3.2 mm stainless steel bead (BioSpec Products). The samples were resuspended in 500 μL of Buffer A (200 mM NaCl (DOT Scientific), 20 mM EDTA (Sigma) and 200 mM Tris·HCl pH 8.0 (Research Products International)), 210 μL 20% SDS (Alfa Aesar) and 500 μL phenol/chloroform/isoamyl alcohol (Invitrogen). Cells were lysed by mechanical disruption with a bead-beater (BioSpec Products) for 3 min twice to prevent overheating. Next, cells were centrifuged for 5 min at 8,000 x g at 4°C, and the supernatant was transferred to a microcentrifuge tube. We added 60 μL 3M sodium acetate (Sigma) and 600 μL isopropanol (LabChem) to the supernatant and incubated on ice for 1 hr. Next, samples were centrifuged for 20 min at 18,000 x g at 4°C. The harvested DNA pellets were washed once with 500 μL of 100% ethanol (Koptec). The remaining trace ethanol in the sample was removed by air drying. Finally, the DNA pellets were then resuspended into 200 μL of AE buffer (Qiagen) left in 4°C overnight to facilitate dissolving. The crude DNA extracts were purified by a Zymo DNA Clean & Concentrator™-5 kit (Zymo Research). The following PCR amplification and NGS sequencing procedures are the same as previously described for treatment of *in vitro* samples.

### *C. difficile* toxin measurements using ELISA

The total toxin concentration (including both TcdA and TcdB) in mice cecal contents were determined by comparison to a standard curve using ELISA (tgcBiomics, Germany). Samples compared for statistical analysis in the same experiment were processed in parallel at the same time using the same batch of ELISA kits to minimize batch-to-batch variations and ensure comparable results.

### *C. difficile* CFU counting from fecal and cecal samples

*C. difficile* selective plates were prepared by autoclaving *C. difficile* agar (Oxoid) and adding taurocholic acid sodium salt hydrate (Sigma, 1 g/L media), D-cycloserine (Sigma, 250 mg/L), and cefoxitin sodium salt (Sigma, 20 mg/L) after the media was cooled to ∼55 °C. Immediately following fecal or cecal collection from mice, around 1 μL of fresh samples were taken using an inoculating loop and mixed with PBS. The samples were then serially diluted (1:10 dilution) using PBS. Six dilutions of each sample were spotted on *C. difficile* selective agar plates. Plates were incubated at 37 °C for 24 h. At this point, colonies were counted in the dilution spot and contained 5-100 colonies. The lower limit of detection for the assay was 20,000 CFU/ mL.

## Resource availability

### Lead contact

Further information and requests for resources and reagents should be directed to and will be fulfilled by the lead contact, Ophelia S. Venturelli.

### Data and code availability

Raw 16S rRNA amplicon sequencing files have been deposited at Zenodo (https://zenodo.org/records/14588768) and are publicly available as of the date of the publication. DOIs are listed in the key resources table. The processed data for all community experiments are available in Github Repository (https://github.com/VenturelliLab/Cdiff_Liu_2025). Processed data with replicates are available in **Table S5**. Scripts for model training and analysis are available in Github Repository (https://github.com/VenturelliLab/Cdiff_Liu_2025). Additional information required to reanalyze the data reported in this paper are available from the lead contact under reasonable request.

## Acknowledgements

We would like to thank all current and past members of the Venturelli lab for their helpful discussions. We would like to thank Dr. Jordy Sulaiman and Dr. Susan Hromada for their advice related to *C. difficile* culturing and experimentation. Research was sponsored by the National Institutes of Health under Grant Number R01AI94497, R01EB030340 and Army Research Office W911NF-19-1-0269.

## Author contributions

Y.L. and O.S.V conceived the research. H.I. and Y.L. performed the *in vitro* monoculture and community experiments. H.I., Y.L., J.T., G.M., K.L. implemented computational modeling. L.L. and D.A.N. performed exo-metabolomics. H.I. and Y.L. performed germ-free mouse experiments. H.I., Y.L., J.T., G.M., K.L., R.B., and S.M.G. analyzed data. H.I., Y.L., and O.S.V. wrote the manuscript and all authors provided feedback on the manuscript. O.S.V. secured funding.

