## Supplementary Information for "Mechanistic and machine learning models design human gut consortia that robustly inhibit *Clostridioides difficile*"

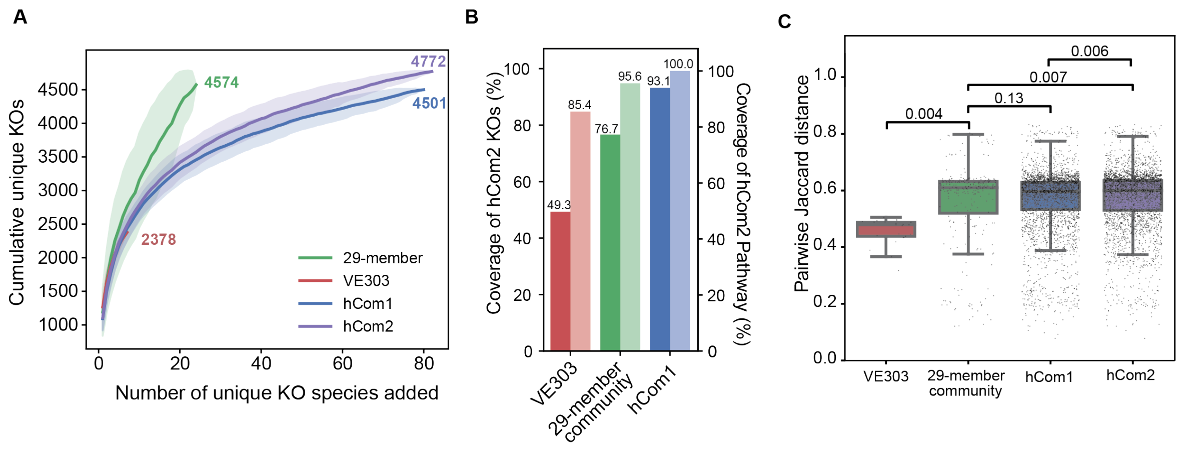


**Supplementary Figure 1. Metabolic functional diversity comparison of synthetic gut communities. (A)** The total number of unique KEGG Orthologs (KOs) accumulation curves generated by iterative random sampling of species (100 permutations). The 29-member community (n = 24 species with KEGG annotations) contained 4,574 unique KOs, compared to 2,378 in VE303 (n = 7), 4,501 in hCom1 (n = 80), and 4,772 in hCom2 (n = 82). **(B)** Distribution of pairwise Jaccard similarity indices computed between all species pairs within each community based on KO profiles. VE303 (mean = 0.545, n = 21 pairs), 29-member community (mean = 0.447, n = 276 pairs), hCom1 (mean = 0.456, n = 3160 pairs), and hCom2 (mean = 0.428, n = 3,321 pairs). Statistical significance was calculated by unpaired t-test. **(C)** Fractional coverage of the hCom2 KO repertoire (left y-axis) and KEGG metabolic pathway (right y-axis) by each community.

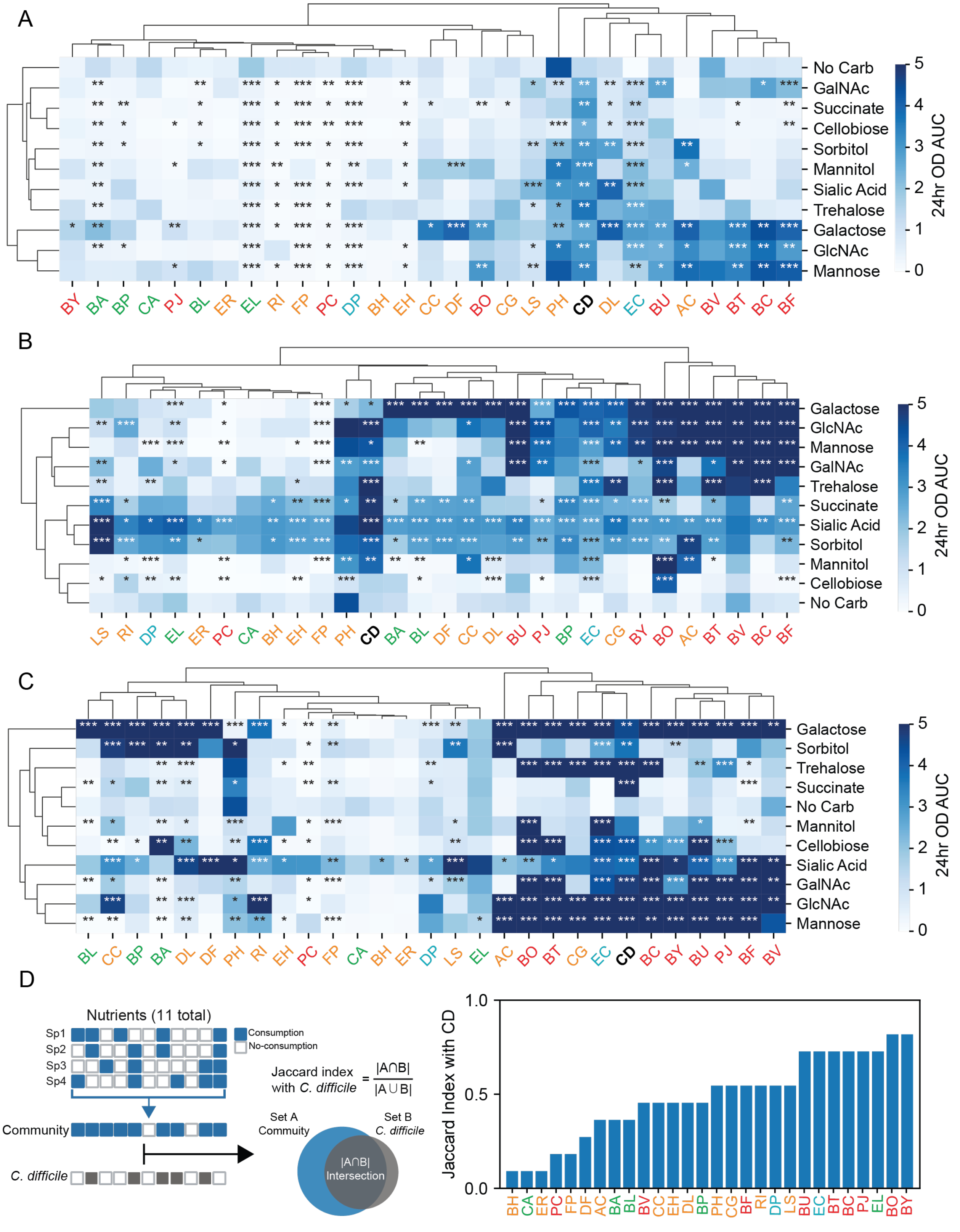


**Supplementary Figure 2. Monoculture growth in single carbohydrate media, related to Figure 1.** (A-C) Clustered heatmap of monoculture fitness of human gut commensals in 1 g/L (A),

5 g/L (B), or 10 g/L (C) single carbohydrate media. The 10 single carbohydrate media are defined media with 1g/L (A), 5g/L (B), or 10g/L (C) of each of the ten carbohydrates from the metabolite design space. A no carbohydrate media condition is included to study the effect of amino acids on species growth. The monoculture fitness is determined by the area under the curve (AUC) of 24 h growth curves. The heatmap is clustered using the Seaborn Clustermap module[1]. Species colors represent phylum. Target species *C. difficile* (CD) is bolded and highlighted in black. Asterisks represent statistical significance compared to No Carb condition: *P < 0.05, **P < 0.01, ***P < 0.001 according to an unpaired t-test. (D) Jaccard index of resource utilization overlap between *C. difficile* and each species in CR-mono model. The Jaccard index is calculated by the number of resources utilized by both *C. difficile* and the species of interest divided by the total number of resources. Resources with a $C_{ij}$ conversion coefficient at least two standard deviations from 0 are classified as resources utilized by a species. Species are ranked by Jaccard index from low to high. Species colors represent phylogeny.

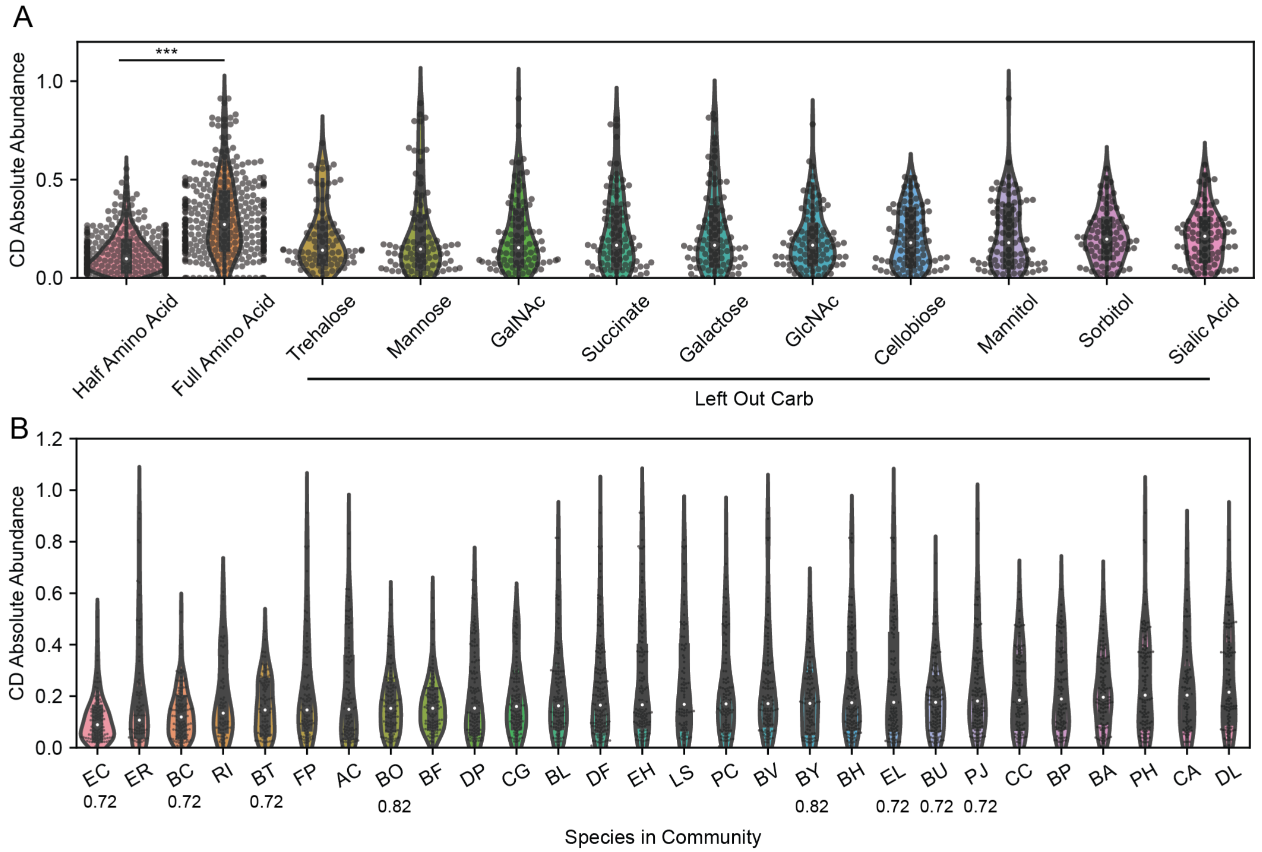


**Supplementary Figure 3. Data from nutrient perturbation experiments in DTL0 and community experiment method validations, related to Figure 2.** (A) Violin plot of *C. difficile* abundance by a resource in media. 20 different media conditions were investigated, including single substrate dropouts at two different amino acid concentrations. Species ODs were sampled at 24 h and were determined by total cell density ($OD_{600}$) and species relative abundance. Species relative abundances are determined using multiplexed 16S rRNA sequencing. Each datapoint is a unique experimental condition averaged over 2-3 replicates. Asterisks represent statistical significance: $*P < 0.05$, $**P < 0.01$, $***P < 0.001$ according to an unpaired t-test. (B) Violin plot of *C. difficile* abundance by species in the inoculum community in DTL0. Each violin represents the distribution of *C. difficile* absolute abundance in experimental conditions where a given species is present in the community. Species ODs were sampled at 24 h and were determined by total cell density ($OD_{600}$) and species relative abundance. Species relative abundances are determined using multiplexed 16S rRNA sequencing. Each datapoint is a unique experimental condition averaged over 2-3 replicates. Species are sorted from low to high median *C. difficile* absolute abundance. The Jaccard index with *C. difficile* from CR-mono model for high overlap species is marked under species names, specifically 0.82 for BO and BY and 0.72 for EC, BC, BT, EL, BU, and PJ.

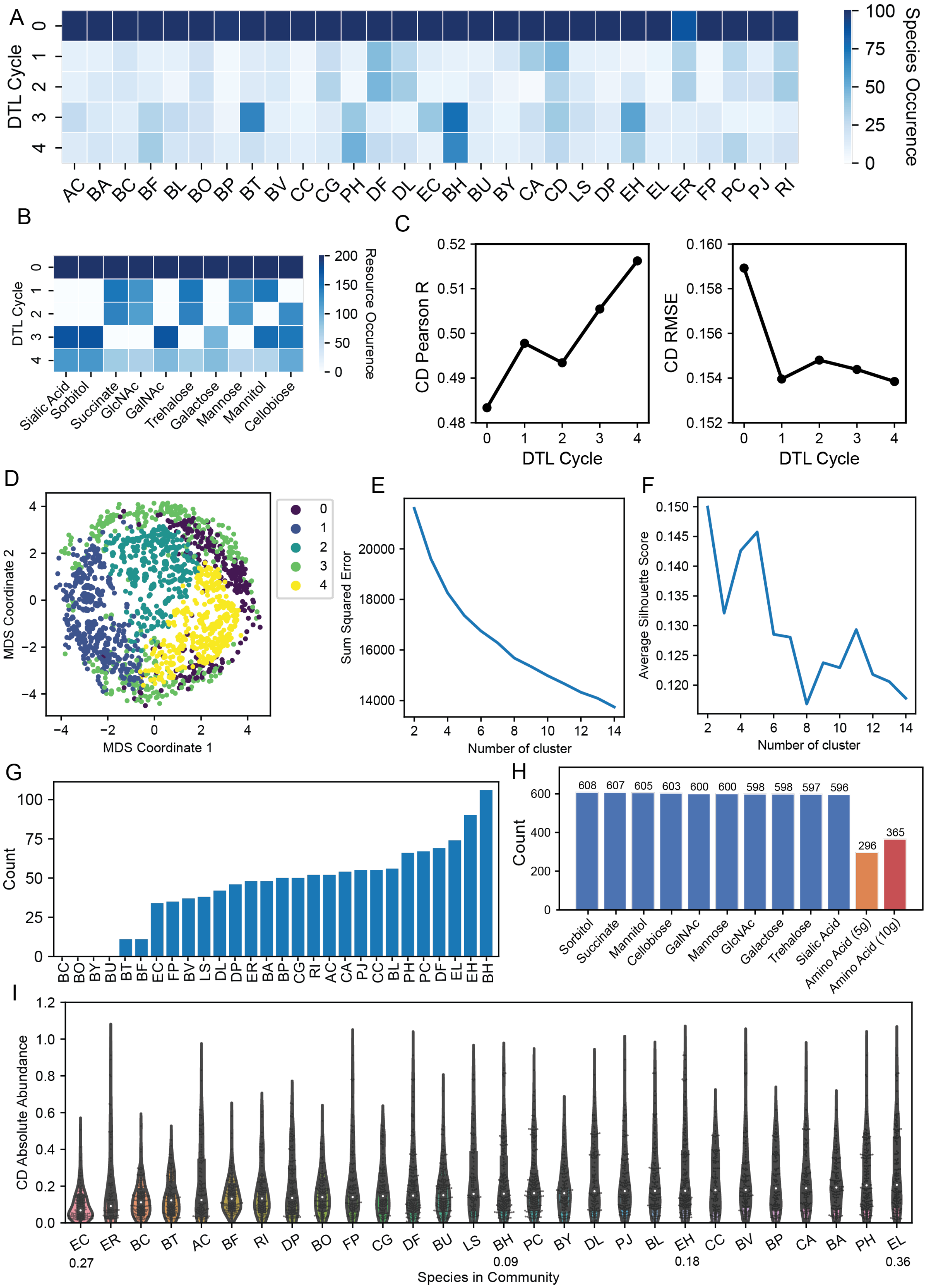

A
B
C
D
E
F
G
H
I
DTL Cycle
Species Occurence
Resource Occurence
CD Pearson R
CD RMSE
MDS Coordinate 1
MDS Coordinate 2
Sum Squared Error
Number of cluster
Average Silhouette Score
Count
CD Absolute Abundance
Species in Community
Sialic Acid
Sorbitol
Succinate
GlcNAc
GalNAc
Trehalose
Galactose
Mannose
Mannitol
Cellobiose
Amino Acid (5g)
Amino Acid (10g)
0.27
0.09
0.18
0.36

**Supplementary Figure 4. Experimental conditions in each DTL cycle and high *C. difficile* abundance cluster, related to Figure 2.** (A), (B) Heatmap of model-selected species (A) and metabolite (B) experimental conditions in each DTL cycle. Occurrence is determined by the count of experimental conditions in each DTL cycle that includes a given species (A) or metabolite (B). All species combinations of monoculture and 2-member, 3-member, and 4-member communities were included in the design algorithm for each DTL cycle. For computational efficiency and feasibility, DTL1, DTL2, DTL3 each included 5 out of 10, 5 out of 10, and 6 out of 10 carbohydrates in the design algorithm, respectively. (C) Consumer resource (CR) model prediction performance of *C. difficile* abundance evaluated by Pearson correlation and root means squared error (RMSE) across DTL cycles. Pearson correlation and RMSE are calculated between predicted and experimental CD abundance in 20-fold cross-validation. (D) Multidimensional scaling (MDS) principal coordinate plots of all DTL and validation community data colored by cluster assignments. KMeans clustering is performed on experimental conditions represented by binary vectors that capture presence or absence of species phylogeny and metabolites, using Scikit-Learn's KMeans function[2]. (E), (F) Plots of sum squared error (E) and average silhouette score (F) are used to select number of clusters. Sum squared error is obtained from the inertia of Scikit-Learn's KMeans function. Average silhouette score is obtained using Scikit-Learn's Silhouette Score function[2]. (G, H) Bar plots of inoculum species counts (G) and metabolite condition counts (H) across experimental conditions in the high *C. difficile* abundance cluster (cluster 4). All DTL data conditions assigned to cluster 4 are used for counting. In (G), each bar is the number of cluster 4 conditions in which a given species is present in the inoculum community. Each bar represents the number of cluster 4 conditions (n = 661) containing a given species (G) or metabolite (H). Carbon sources are scored as present/absent and amino acids are separated by supplemented amount (5 g/L and 10 g/L). (I) Violin plot of *C. difficile* abundance by species in the inoculum community in all DTL data. Each violin represents the distribution of *C. difficile* absolute abundance in experimental conditions where a given species is present in the community. Species ODs were sampled at 24 h and were determined by total cell density (OD600) and species relative abundance. Species relative abundances are determined using multiplexed 16S rRNA sequencing. Species are sorted from low to high median *C. difficile* absolute abundance.

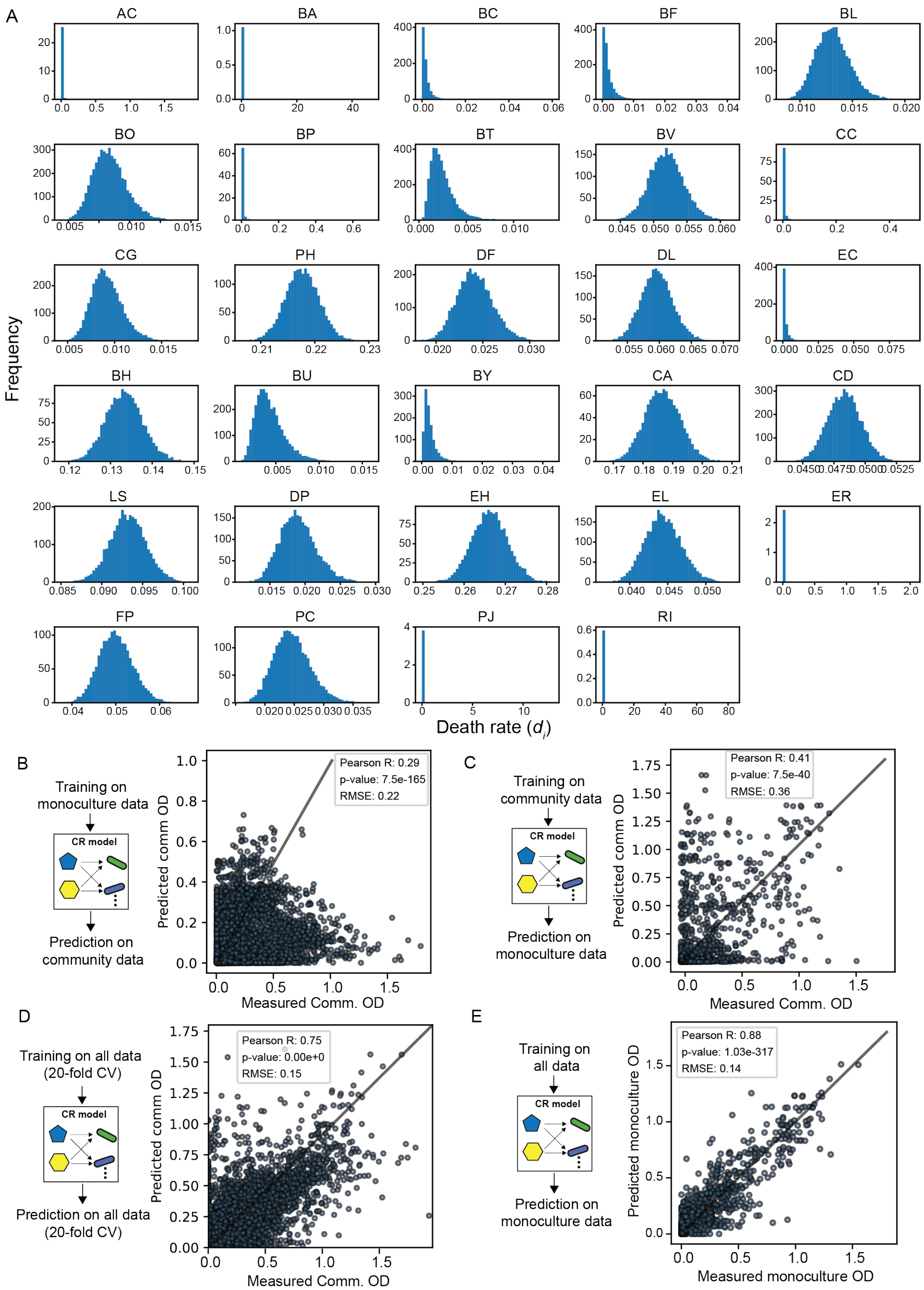


**Supplementary Figure 5. CR model parameters and predictive performance.** (A) CR-comm model predicted death rates for each species. The CR-comm model is trained on all DTL data including both monoculture and community growth dynamics. The x-axis represents the death rate parameter ($d_i$) and the y-axis represents the frequency across mono and community conditions. (B–E) Prediction performance of the CR model under different training and evaluation schemes: (B) trained on monoculture data only and evaluated on all community conditions, (C) trained on community data only and evaluated on all monoculture conditions, (D) 20-fold cross-validation using all data, and (E) fit to monoculture data when trained on all monoculture and community data.

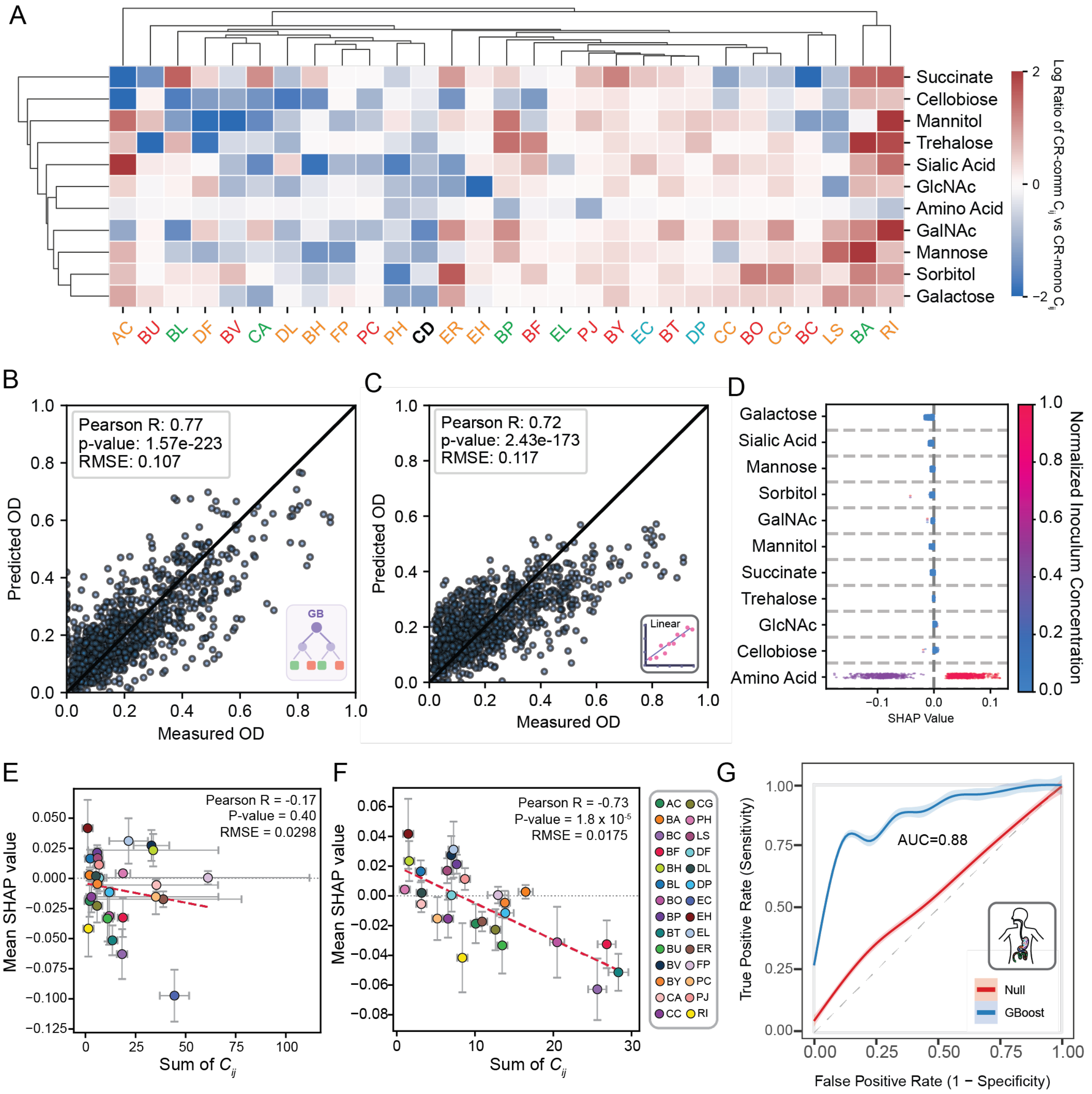

**Supplementary Figure 6.** (A) Clustered heatmap of the log ratio of conversion coefficients, $C_{ij}$, between the CR-comm model and the CR-mono model. Colors indicate whether the conversion coefficient is higher in the CR-comm model (red) or the CR-mono model (blue). (B and C) Scatterplots for measured versus predicted 24 h *C. difficile* OD from the gradient boosting (GB) model (B) and a modified linear regression model (C). 20-fold cross-validation was performed on monoculture and community growth data from all DTL cycles. Prediction performance was evaluated by Pearson correlation coefficient and RMSE between measured and predicted 24 h *C. difficile* OD. For the linear regression model, one outlier at (0.38, −47,935.51) is not depicted. (D) Distribution of SHAP values for metabolite features in the GB model. SHAP values were calculated across all experimental conditions in which each metabolite is present. Each data point represents the feature contribution in a single experimental condition. Features are sorted by mean SHAP value from negative to positive. Colors indicate the normalized inoculum metabolite concentration, calculated by dividing the experimental inoculum concentration by the maximum inoculum concentration across the entire dataset. (E and F) Scatter plots of GB-model SHAP value versus summed $C_{ij}$ value across strains. Each dot represents an individual strain. The y-axis is the mean GB-model SHAP value for that strain, computed over the communities in which the strain is present (OD > 0). Error bars denote ±1 standard deviation of those SHAP values across communities. The x-axis is the sum of the strain's positive consumption coefficients ($C_{ij} > 0$) across all 11 nutrients in the CR model. Error bars denote ±1 standard deviation of the summed $C_{ij}$, propagated from the posterior parameter uncertainty of the CR models: $C_{ij}$ parameters from the monoculture-fit model (CR-mono) (E) and the community-fit model (CR-comm) excluding *E. coli* (F). The squared posterior standard deviations of the contributing coefficients were summed and the square root taken, i.e. $\sigma(summed\ C_{ij}) = \sqrt{\sum_i \sigma^2{}_{ij}}$. The dashed red line is the linear least-squares fit. Colors indicate species identity. (G) Receiver operating characteristic (ROC) curve of a GB model trained on human gut microbiome data to predict *C. difficile* presence. The area under the ROC curve (AUC) is annotated in the figure.

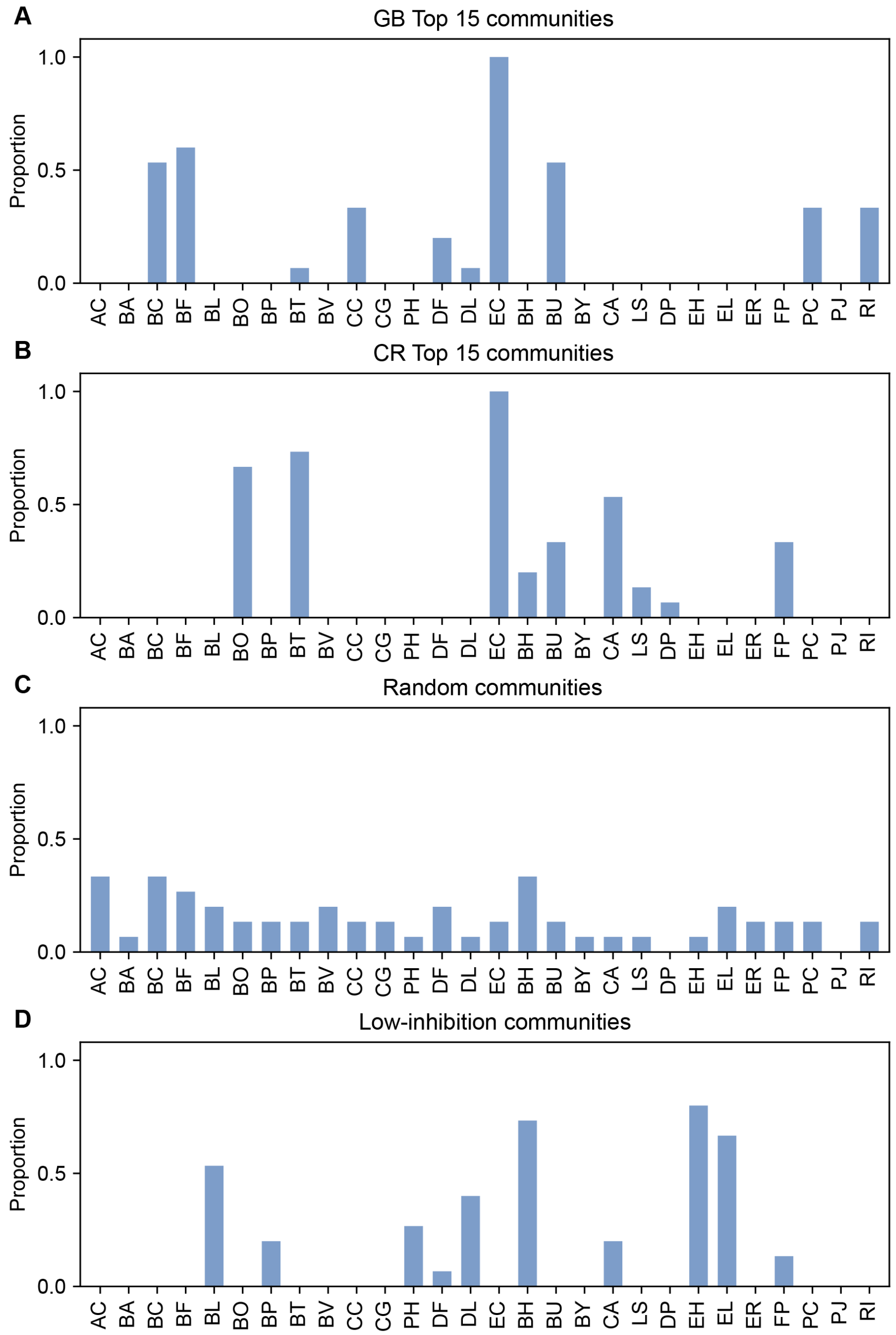


**Supplementary Figure 7. Species counts in *in vitro* validation communities, related to Figure 4.** (A-D) Species counts in 15 optimized communities in CR-opt (A), GB-opt (B), random (C), or low-inhibition set designed by CR and GB model (D). The 15 optimized communities designed by CR or GB model were designed by minimizing the maximum 24 h *C. difficile* OD across 22 *in silico* media conditions. The random communities were designed by the Numpy Random package[3]. The low-inhibition communities were designed by maximizing the minimum 24 h *C. difficile* OD across 22 *in silico* media conditions in both CR and GB model predictions.

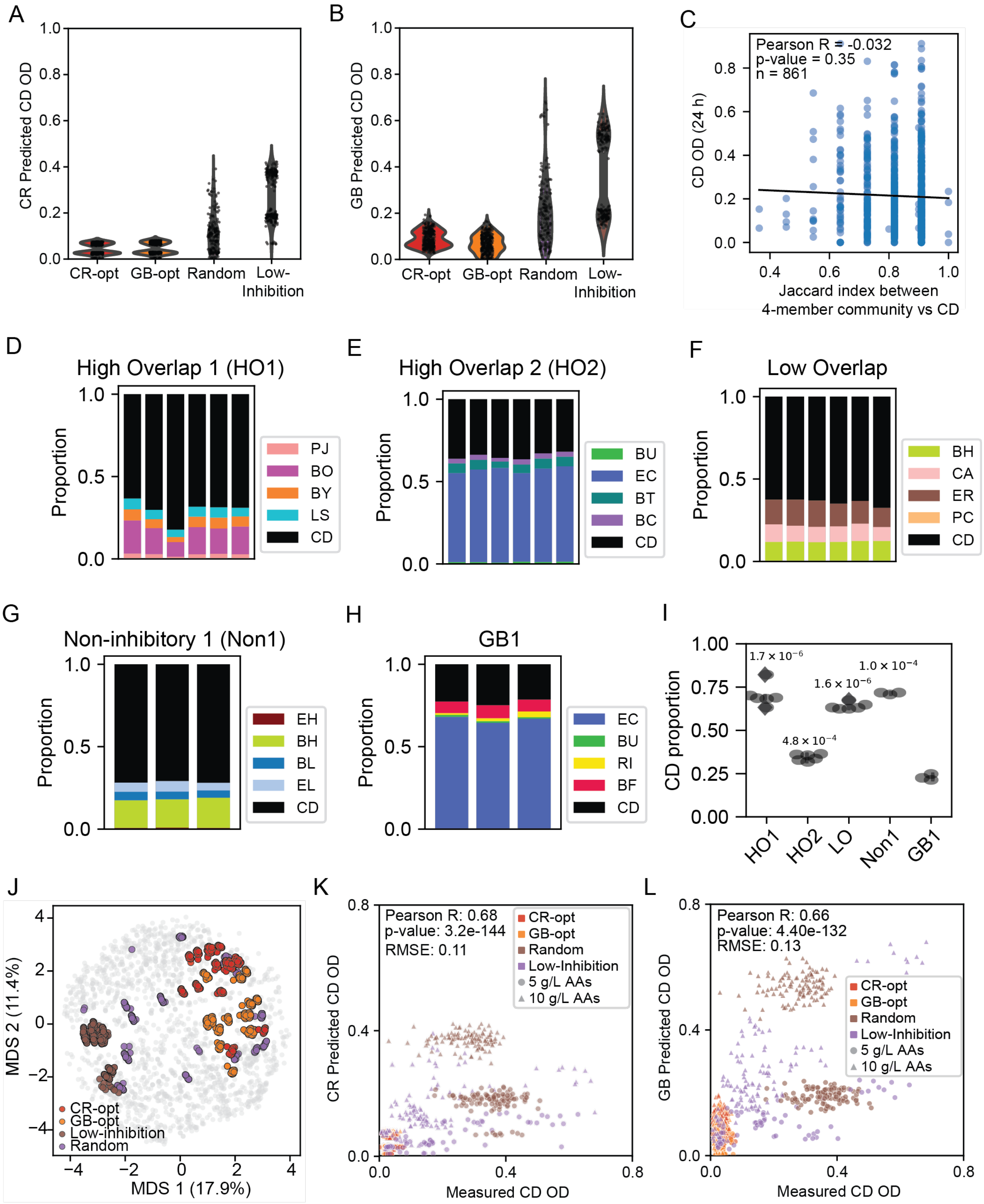


**Supplementary Figure 8. Model prediction performances of *in vitro* validation communities and species abundance analysis of the communities designed on the niche-overlap data, related to Figure 4.** (A), (B) CR (A) and GB (B) model predicted 24 h *C. difficile* OD distribution of CR-opt (red, 15 communities optimized by the CR-comm model for low predicted *C. difficile*

$OD_{600}$), GB-opt (orange, 15 communities optimized by the GB model), Random (purple, 15 random-composition controls), and Low-Inhibition groups (brown, 15 non-inhibitory communities with high predicted *C. difficile* $OD_{600}$ by both CR-comm and GB models). Distributions represent predicted 24 h *C. difficile* OD across 22 *in silico* media conditions. (C) Scatter plot of Jaccard index of resource utilization overlap between each 4-member community and *C. difficile* plotted against measured *C. difficile* $OD_{600}$ at 24 h (n = 861). The black line represents the ordinary least-squares linear fit. The Jaccard index for each community was calculated as the fraction of resources utilized by both the community and *C. difficile*, based on the species-level resource utilization profiles from the CR-mono model (Figure S2D). (D-H) Stacked bar plots show the relative abundances of species in High-overlap 1 (HO1) (D), High-overlap 2 (HO2) (E), Low-Overlap (LO) (F), Non-inhibitory 1 (Non1) (G), and GB1 (H) communities after 24h of co-incubation with CD, measured by 16S rRNA sequencing. The HO1 and HO2 communities were designed by high niche overlap with *C. difficile* while the LO community was designed by low niche overlap with *C. difficile*. The degree of niche overlap was calculated using the values from CR-mono model. The HO1 community consists of BO, BY, CS, and PJ; HO2 includes BC, BT, BU, and EC; and the LO community includes BH, CA, ER, and PC. (I) Box and scatter plots show the absolute abundance of CD across the different communities. Statistical significance was calculated by unpaired t-test respective "GB1" condition. (J) Multidimensional scaling (MDS) projection of the communities with resources design space (see Methods). Colored points denote validation communities by design group. (K and L) Scatter plots of measured 24 h *C. difficile* OD versus predicted 24 h *C. difficile* OD of CR-comm model (K) and GB model (L) for *in vitro* validation. Colors and markers indicate validation group identity and amino acid levels, respectively. Prediction performances are evaluated by Pearson correlation and RMSE between measured and predicted 24 h *C. difficile* OD. The prediction Pearson correlation for Random group by CR is 0.42, with P-value of $1.79*10^{-9}$. The prediction Pearson correlation for Random group by GB is 0.58, with P-value of $2.30*10^{-18}$. The percent of correct prediction of CD OD either above or below 0.01 (5 times the initial OD of 0.002) for each group by the CR model is 92% (CR-opt), 95% (GB-opt), 95% (Random), and 100% (Low-Inhibition). The percent of correct prediction of CD OD either above or below 0.01 (5 times the initial OD of 0.002) for each group by the GB model is 97% (CR-opt), 89% (GB-opt), 98% (Random), and 100% (Low-Inhibition).

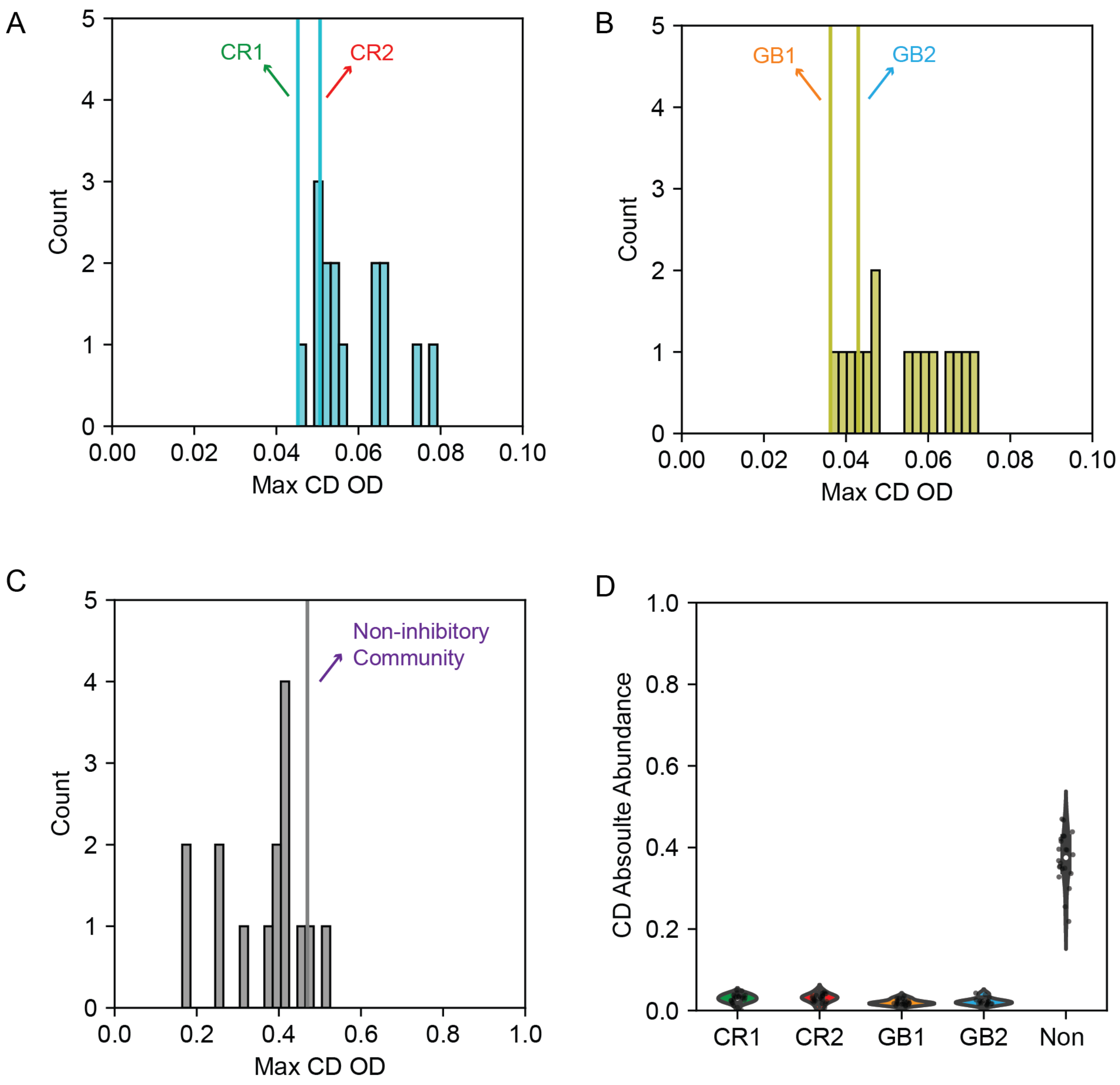


**Supplementary Figure 9. Predictions and measured 24 h *C. difficile* OD distributions for exo-metabolomic experiment communities.** (A-C) Distribution of maximum 24 h *C. difficile* OD in CR-opt communities (A), GB-opt communities (B), and low-inhibition communities designed by CR-comm and GB models (C). The maximum *C. difficile* OD for each community is determined from the maximum 24 h *C. difficile* OD *in vitro* validations across 22 media conditions. The maximum *C. difficile* OD for communities selected for metabolomics are marked by vertical lines (CR1, CR2, GB1, GB2, and Non-inhibitory community 1). (D) Measured *in vitro* 24 h *C. difficile* OD distribution of communities selected for exo-metabolomics experiment across 22 media conditions. Each datapoint represents average over 3 biological replicates.

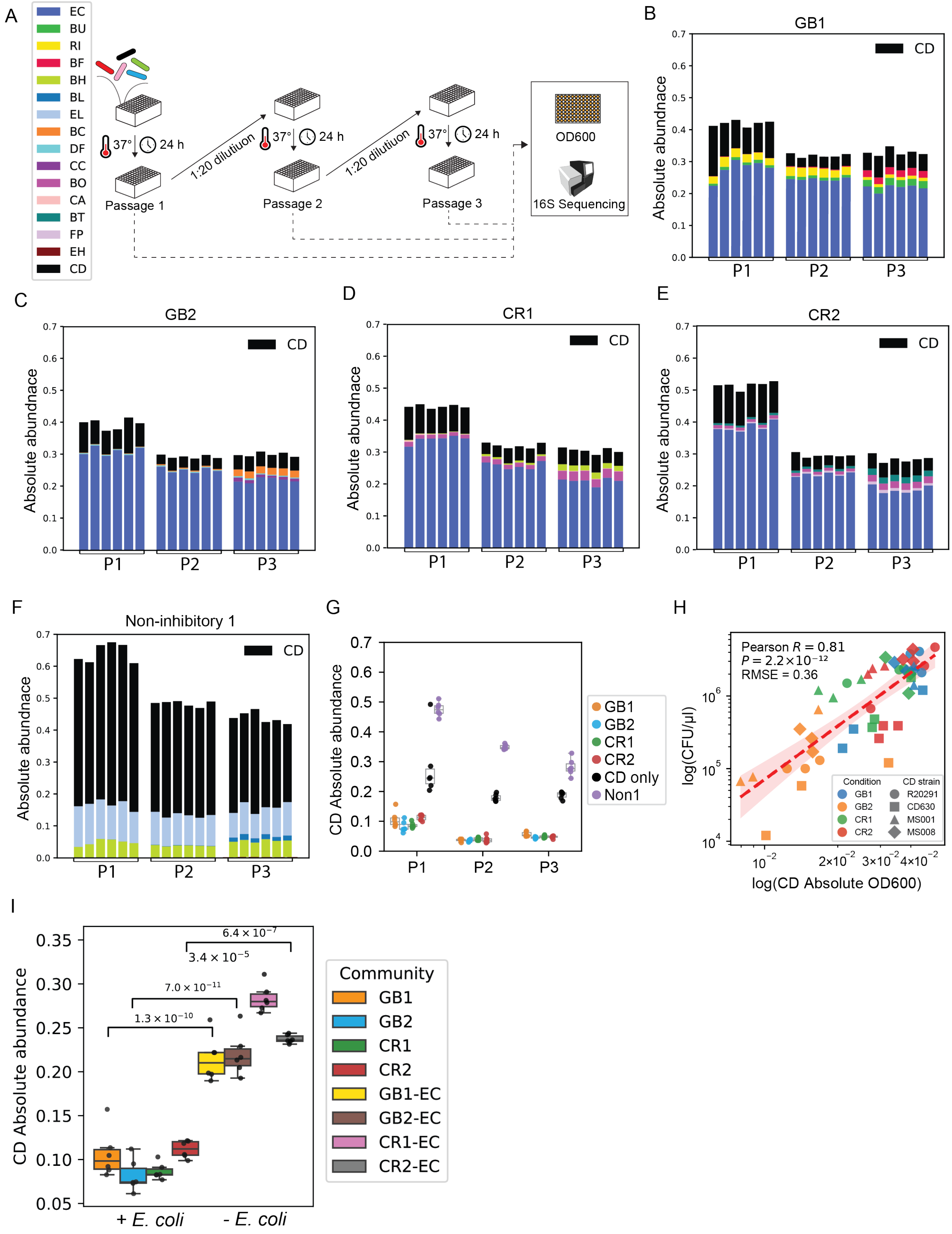
A
EC
BU
RI
BF
BH
BL
EL
BC
DF
CC
BO
CA
BT
FP
EH
CD
37°
24 h
Passage 1
1:20 dilutiuon
Passage 2
Passage 3
OD600
16S Sequencing
B
GB1
C
GB2
D
CR1
E
CR2
F
Non-inhibitory 1
Absolute abundnace
P1
P2
P3
G
CD Absolute abundance
GB1
GB2
CR1
CR2
CD only
Non1
H
Pearson R = 0.81
P = 2.2×10⁻¹²
RMSE = 0.36
log(CFU/μl)
log(CD Absolute OD600)
Condition
CD strain
R20291
CD630
MS001
MS008
I
CD Absolute abundance
6.4 × 10⁻⁷
3.4 × 10⁻⁵
7.0 × 10⁻¹¹
1.3 × 10⁻¹⁰
+ E. coli
- E. coli
Community
GB1
GB2
CR1
CR2
GB1-EC
GB2-EC
CR1-EC
CR2-EC

**Supplementary Figure 10. *C. difficile* inhibition performance of model-optimized communities.** (A) Schematic of the passaging experiments. Synthetic communities were passaged every 24 h with a 1:20 dilution. Absolute abundance of each strain was determined by multiplying its relative abundance by the total cell density ($OD_{600}$) measurement. (B–G) Stacked bar plots showing absolute species abundances in the selected representative communities: GB1 (B), GB2 (C), CR1 (D), CR2 (E), and non-inhibitory 1 (F). Bar colors indicate species identity. (G) Scatter plots showing the absolute abundance of *C. difficile* across different communities. Statistical significance was assessed by ANOVA, with *P*-values annotated in the plots. (H) Scatter plot of absolute abundance versus viable *C. difficile* cell counts. Data were collected from four synthetic communities (GB1, GB2, CR1, and CR2) containing *C. difficile* strains (R20291, CD630, MS001, and MS008). TCCFA medium was used to selectively culture viable *C. difficile* cells from the communities[4]. A linear regression fit (red dashed line) with 95% confidence interval (shaded area) indicates a strong positive correlation between absolute abundance and CFU (Pearson R = 0.81, $P = 2.2 \times 10^{-12}$, RMSE = 0.36). Each point represents a biological replicate. (I) Bar and scatter plots showing absolute abundance of *C. difficile* across the different communities. Statistical significance between conditions with and without *E. coli* was assessed by unpaired t-test. P-values are indicated on the plot. The color of bar indicates the community condition.

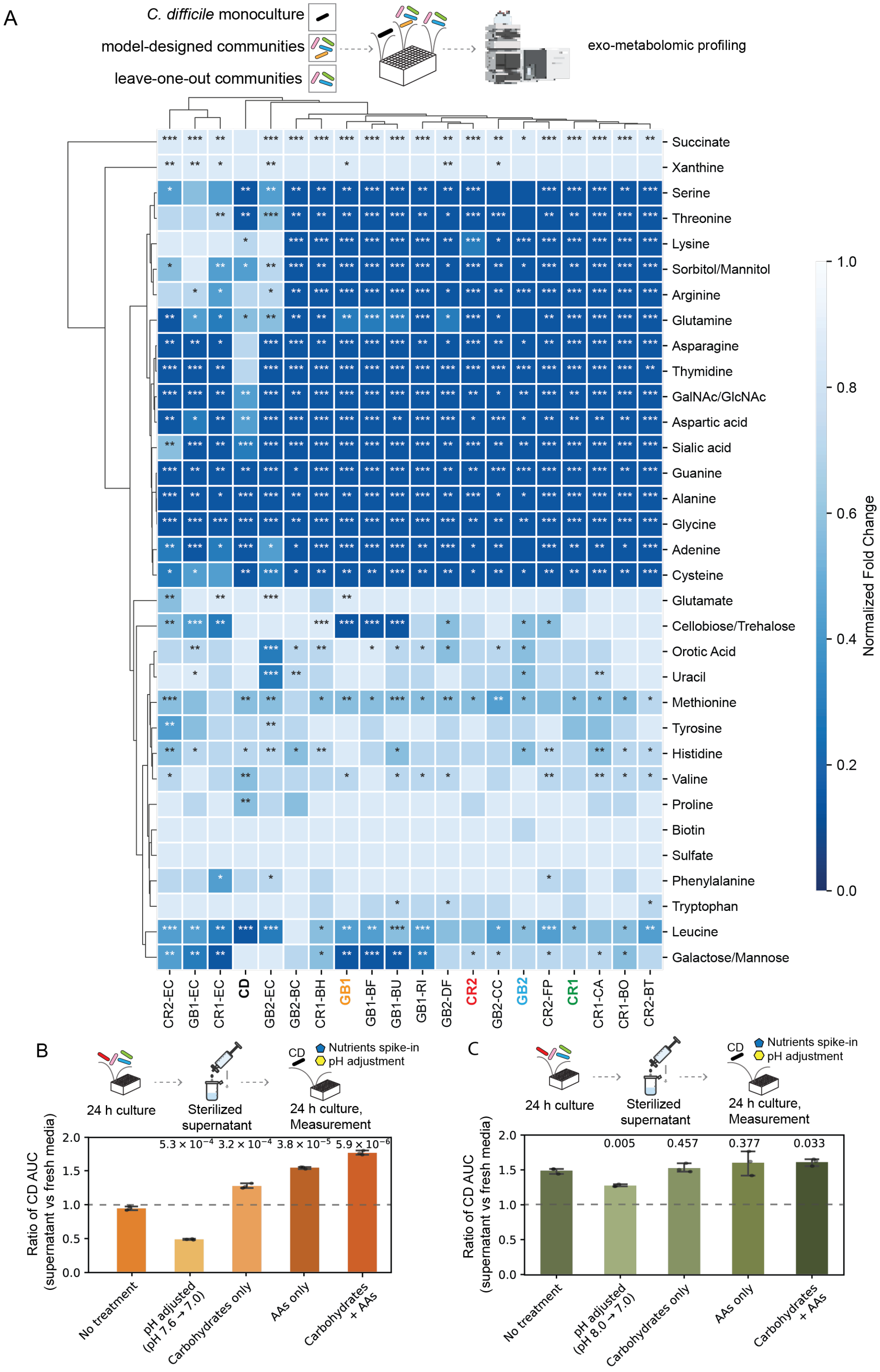
A
C. difficile monoculture
model-designed communities
leave-one-out communities
exo-metabolomic profiling
Succinate
Xanthine
Serine
Threonine
Lysine
Sorbitol/Mannitol
Arginine
Glutamine
Asparagine
Thymidine
GalNAc/GlcNAc
Aspartic acid
Sialic acid
Guanine
Alanine
Glycine
Adenine
Cysteine
Glutamate
Cellobiose/Trehalose
Orotic Acid
Uracil
Methionine
Tyrosine
Histidine
Valine
Proline
Biotin
Sulfate
Phenylalanine
Tryptophan
Leucine
Galactose/Mannose
CR2-EC
GB1-EC
CR1-EC
CD
GB2-EC
GB2-BC
CR1-BH
GB1
GB1-BF
GB1-BU
GB1-RI
GB2-DF
CR2
GB2-CC
GB2
CR2-FP
CR1
CR1-CA
CR1-BO
CR2-BT
Normalized Fold Change
1.0
0.8
0.6
0.4
0.2
0.0
B
24 h culture
Sterilized supernatant
CD
Nutrients spike-in
pH adjustment
24 h culture, Measurement
Ratio of CD AUC (supernatant vs fresh media)
$5.3 \times 10^{-4}$
$3.2 \times 10^{-4}$
$3.8 \times 10^{-5}$
$5.9 \times 10^{-6}$
No treatment
pH adjusted (pH 7.6 → 7.0)
Carbohydrates only
AAs only
Carbohydrates + AAs
C
24 h culture
Sterilized supernatant
CD
Nutrients spike-in
pH adjustment
24 h culture, Measurement
Ratio of CD AUC (supernatant vs fresh media)
0.005
0.457
0.377
0.033
No treatment
pH adjusted (pH 8.0 → 7.0)
Carbohydrates only
AAs only
Carbohydrates + AAs

**Supplementary Figure 11. Metabolite profiling of model-designed communities and supernatant-mediated *C. difficile* growth inhibition.** (A) Clustered heatmap of normalized metabolite fold changes in *C. difficile* monoculture, model-designed communities, and leave-one-out communities. Model-designed communities include two four-member communities designed by GB model (GB1 and GB2) and two four-member communities designed by CR model (CR1 and CR2). Leave-one-species-out communities are three-member communities based on model-designed communities with one species removed. Species were cultured in media containing 1 g/L of all 10 target carbohydrates with 10 g/L amino acids. Supernatants were harvested at the start of culture (Time 0) and after 24 h of growth (Time 24) for measurement of metabolite concentrations by LC-MS. Absolute fold change of each metabolite was determined by dividing the AUC in the Time 24 sample by that in the Time 0 sample. Colors represent average normalized fold change over 3 biological replicates; darker blue represents lower normalized fold change and higher consumption. Asterisks indicate statistical significance of unpaired t-test: *$P < 0.05$, **$P < 0.01$, ***$P < 0.001$, ns = not significant. (B-C) *C. difficile* growth AUC in GB1 supernatants (B) and in non-inhibitory community 1 supernatants (C). The supernatants were prepared by filter sterilization after 24 h of community culture in fresh defined media containing 1 g/L of all 10 target carbohydrates with 10 g/L amino acids, and pH was adjusted to 7.0 by titration with 1 M HCl. Nutrient spike-ins were performed by adding carbohydrates or amino acids at 1× concentrations equivalent to those in fresh defined media. *C. difficile* AUC values were normalized to that of *C. difficile* grown in fresh defined media. Statistical significance was calculated by unpaired t-test respective "No treatment" condition.

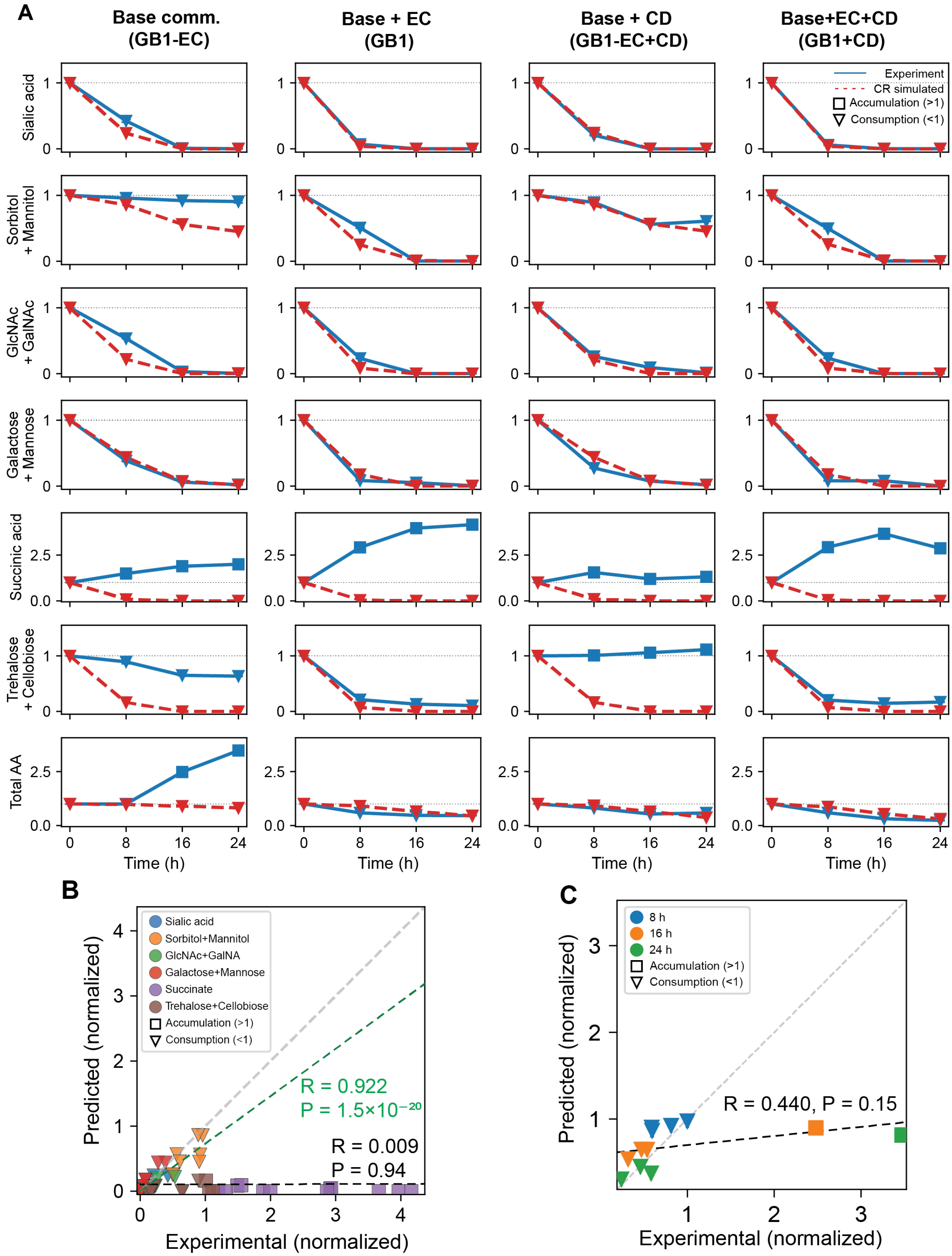
A
Base comm. (GB1-EC)
Base + EC (GB1)
Base + CD (GB1-EC+CD)
Base+EC+CD (GB1+CD)
Experiment
CR simulated
Accumulation (>1)
Consumption (<1)
Sialic acid
Sorbitol + Mannitol
GlcNAc + GalNAc
Galactose + Mannose
Succinic acid
Trehalose + Cellobiose
Total AA
Time (h)
B
Sialic acid
Sorbitol+Mannitol
GlcNAc+GalNA
Galactose+Mannose
Succinate
Trehalose+Cellobiose
Accumulation (>1)
Consumption (<1)
R = 0.922
P = 1.5×10⁻²⁰
R = 0.009
P = 0.94
Predicted (normalized)
Experimental (normalized)
C
8 h
16 h
24 h
Accumulation (>1)
Consumption (<1)
R = 0.440, P = 0.15
Predicted (normalized)
Experimental (normalized)

**Supplementary Figure 12. Comparison of CR-comm model predictions with exo-metabolomics measurements.** (A) Time-course profiles of individual metabolite concentrations as predicted by CR-comm model simulation (red, dashed) and measured experimentally (blue, solid) across four communities (biological replicates, n=2) : Base, Base+EC, Base+CD, and Base+EC+CD. Rows represent individual metabolites: sialic acid, sorbitol/mannitol, GlcNAc/GalNAc, galactose/mannose, succinate, trehalose/cellobiose, and total amino acids. Concentrations are normalized to initial condition (g/L, t = 0 h, y-axis). Total amino acid concentrations were calculated by summing the concentrations of all individual amino acids and normalizing to the initial condition. (B and C) Scatter plots of CR model simulation versus experimentally measured normalized metabolite concentrations across four communities. (B) Carbohydrate metabolites, colored by metabolite. The black dashed line, Pearson correlation coefficient, and associated P-value are computed across all carbohydrates. The green dashed line and corresponding statistics are computed after excluding succinate and trehalose/cellobiose, which exhibit net accumulation due to microbial production. (C) Total amino acid concentration, colored by time point. The black dashed line, Pearson correlation coefficient, and associated P-value are computed across all time points and communities. Marker shape indicates the direction of change relative to the initial concentration at each time point: squares denote net accumulation (normalized value > 1) and downward triangles denote consumption (normalized value < 1). The gray dashed line indicates the identity line (y = x).

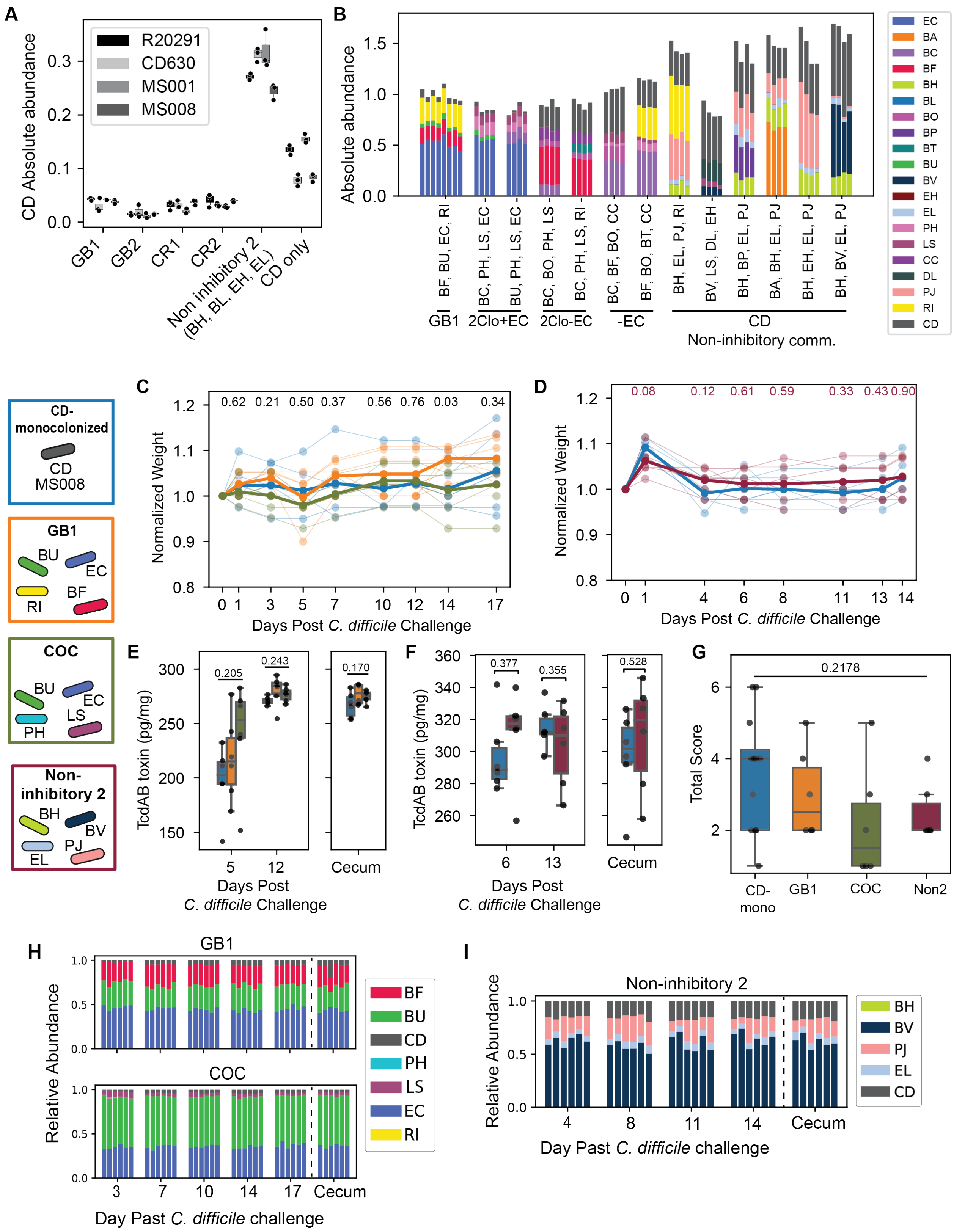

A
CD Absolute abundance
R20291
CD630
MS001
MS008
GB1
GB2
CR1
CR2
Non inhibitory 2 (BH, BL, EH, EL)
CD only
B
Absolute abundance
BF, BU, EC, RI
BC, PH, LS, EC
BU, PH, LS, EC
BC, BO, PH, LS
BC, PH, LS, RI
BC, BF, BO, CC
BF, BO, BT, CC
BH, EL, PJ, RI
BV, LS, DL, EH
BH, BP, EL, PJ
BA, BH, EL, PJ
BH, EH, EL, PJ
BH, BV, EL, PJ
GB1
2Clo+EC
2Clo-EC
-EC
CD
Non-inhibitory comm.
EC
BA
BC
BF
BH
BL
BO
BP
BT
BU
BV
EH
EL
PH
LS
CC
DL
PJ
RI
CD
CD-monocolonized
CD MS008
GB1
BU
EC
RI
BF
COC
BU
EC
PH
LS
Non-inhibitory 2
BH
BV
EL
PJ
C
Normalized Weight
0.62 0.21 0.50 0.37 0.56 0.76 0.03 0.34
Days Post C. difficile Challenge
D
Normalized Weight
0.08 0.12 0.61 0.59 0.33 0.43 0.90
Days Post C. difficile Challenge
E
TcdAB toxin (pg/mg)
0.205
0.243
0.170
Days Post C. difficile Challenge
Cecum
F
TcdAB toxin (pg/mg)
0.377
0.355
0.528
Days Post C. difficile Challenge
Cecum
G
Total Score
0.2178
CD-mono
GB1
COC
Non2
H
GB1
COC
Relative Abundance
BF
BU
CD
PH
LS
EC
RI
Day Past C. difficile challenge
Cecum
I
Non-inhibitory 2
Relative Abundance
BH
BV
PJ
EL
CD
Day Past C. difficile challenge
Cecum

**Supplementary Figure 13. Supplementary data for murine experiment with CD-monocolonized, GB1, COC, and non-inhibitory community 2.** (A) *C. difficile* absolute abundance in model-designed communities across different *C. difficile* strains. Colors indicate *C. difficile* strains. *C. difficile* absolute abundance was calculated by multiplying relative abundance and total cell density ($OD_{600}$) measured after 24 h culture in media containing all 10 target carbohydrates (1 g/L each) with 10 g/L amino acids. Markers represent individual biological replicates (n = 3). (B) Absolute abundance of species composing the model-designed communities. Absolute abundance was calculated after 24 h of culture in media containing all 10 target carbohydrates (1 g/L each) with 10 g/L amino acids. Markers represent individual biological replicates (n = 8 for GB1, n = 4 for other conditions). (C and D) Normalized mouse weight after *C. difficile* challenge. Colors represent experimental groups. Transparent lines represent biological replicates (each mouse) and solid lines represent the average across all biological replicates (n = 6 per group). (E and F) *C. difficile* TcdAB concentration in fecal and cecal samples, measured by ELISA. Colors represent experimental groups and markers represent individual biological replicates (n = 6 per group). (G) Histopathology total score of colon tissue samples from GB1, COC, and non-inhibitory 2 (Non2) communities. Statistical significance was assessed by ANOVA or unpaired t-test. P-values are annotated on the plots. (H and I) Stacked barplots of species relative abundances in model-guided communities. (H) GB1 and COC. (I) Non-inhibitory community 2. Bar colors indicate species identity. Each bar represents an individual biological replicate.

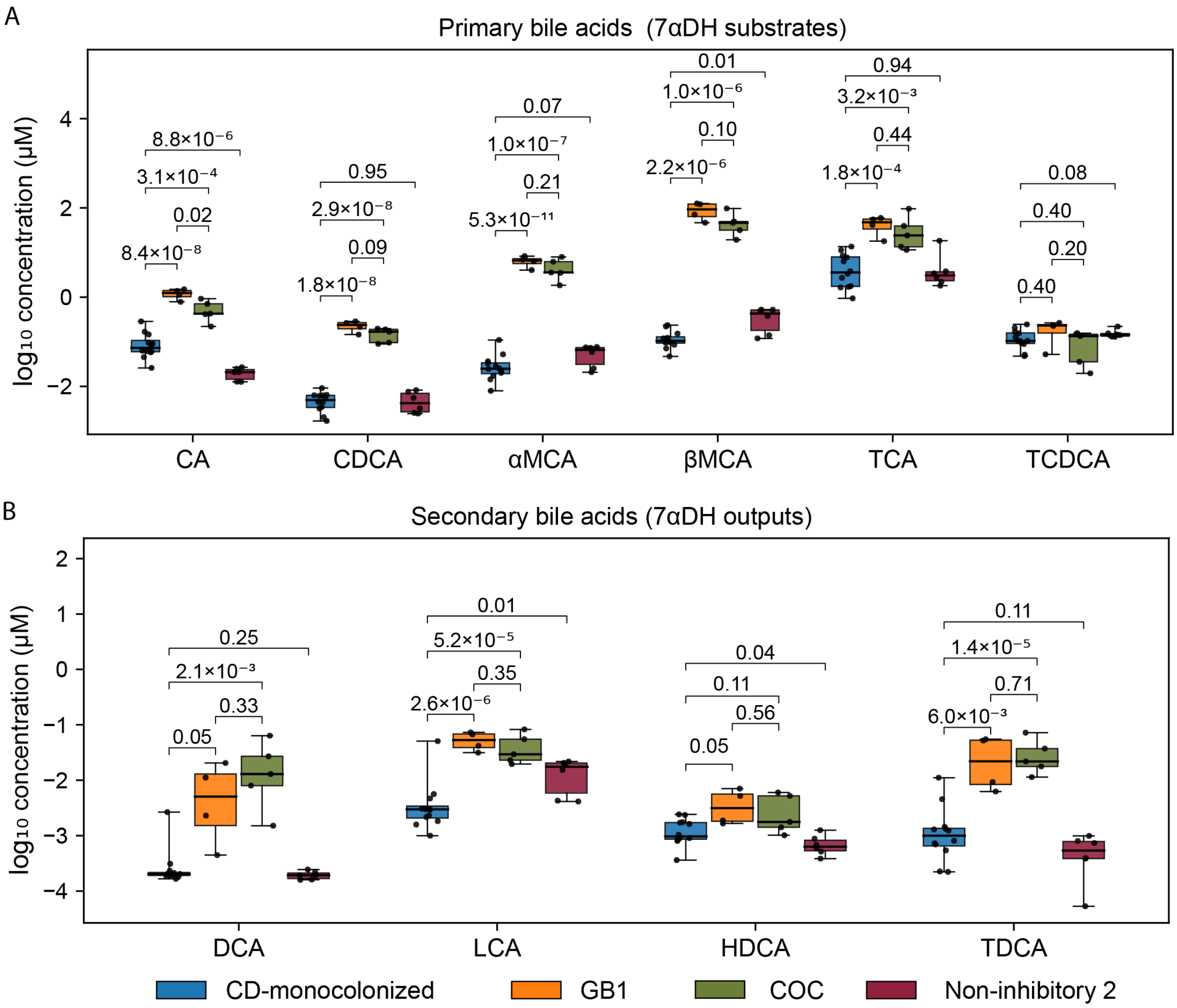


**Supplementary Figure 14. The boxplots of bile acid concentrations in cecal samples from the murine experiment with CD-monocolonized, GB1, COC, and non-inhibitory community 2.** Bile acids were measured in cecal contents collected at the endpoint of the murine experiments (CD-monocolonized, n = 12, pooled from the two mouse experiments; GB1, n = 4; COC, n = 5; Non-inhibitory 2, n = 6). **(A)** Primary bile acids (substrates of 7α-dehydroxylation): cholic acid (CA), chenodeoxycholic acid (CDCA), α-muricholic acid (αMCA), β-muricholic acid (βMCA), taurocholic acid (TCA), and taurochenodeoxycholic acid (TCDCA). **(B)** Secondary bile acids (7α-dehydroxylation products): deoxycholic acid (DCA), lithocholic acid (LCA), hyodeoxycholic acid (HDCA), and taurodeoxycholic acid (TDCA). In each box plot, the box denotes the median and interquartile range, whiskers span the full data range, and individual samples are overlaid as dots colored by condition. Statistical significance was assessed by unpaired t-tests and P-values are annotated in the plots.

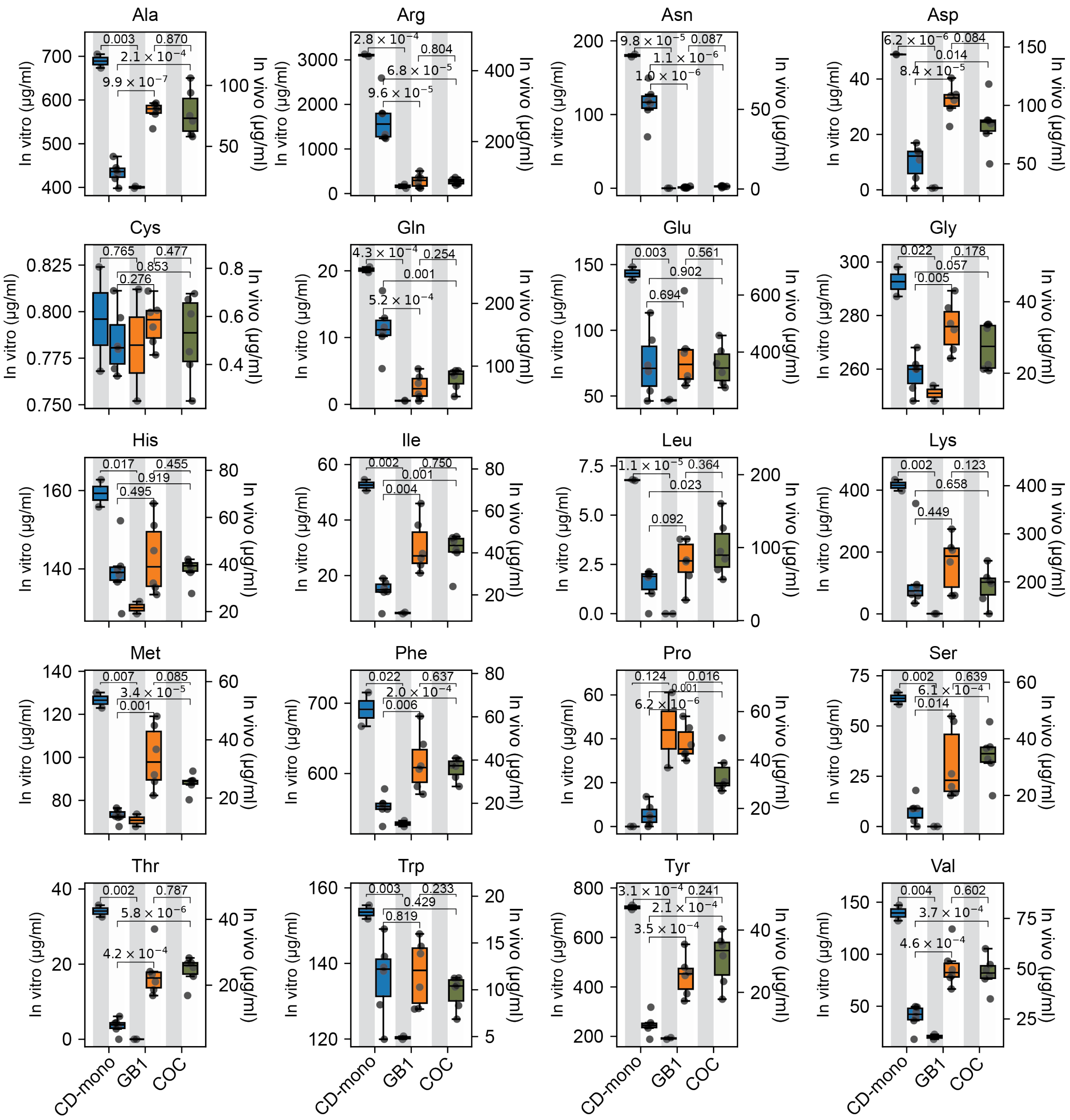


**Supplementary Figure 15. The boxplots of exo-metabolomic analysis of cecal samples from murine experiments with CD-monocolonized, GB1, and COC.** For each condition, two bars are presented: the left bar represents *in vitro* results (gray background) and corresponds to the left y-axis, and the right bar represents *in vivo* results (white background) and corresponds to the right y-axis. *In vitro* samples were prepared from cultures in media containing 1 g/L of all 10 target carbohydrates with 10 g/L amino acids after 16 h (n = 2). *In vivo* samples were collected from cecal contents at the endpoint of the murine experiment (n = 6). Statistical significance was assessed by unpaired t-tests and P-values are annotated in the plots.
